# Surveillance of *in situ* tumor arrays reveals early environmental control of cancer immunity

**DOI:** 10.1101/2021.05.27.445482

**Authors:** Guadalupe Ortiz-Muñoz, Markus Brown, Catherine B. Carbone, Joaquin Pechuan-Jorge, Alex T. Ritter, Darya Orlova, Shreya R. Mantri, Angela Yang, Jonas Doerr, Sanjeev Mariathasan, Yulei Wang, Shannon Turley, Carlos Gomez-Roca, Carlos Eduardo de Andrea, David Spigel, Thomas Wu, Zora Modrusan, Richard Price, Ira Mellman, Christine Moussion

## Abstract

The immune phenotype of a tumor is a key predictor of its response to immunotherapy^1–4^. Patients who respond to immune checkpoint blockade generally present with tumors that are infiltrated by activated T cells, a tumor-immune phenotype referred to as ‘immune inflamed’^5–7^. However, not all immune inflamed tumors respond to therapy, and in addition the majority of patients presents with tumors that lack T cells (‘immune desert’) or that exclude T cells in the periphery of the tumor islet (‘immune excluded’)^8^. Despite the importance of these tumor-immune phenotypes in patients, little is known about their development, heterogeneity or dynamics due to an inability to model these features pre-clinically.

Here, we describe an approach designated STAMP (*s*kin *t*umor *a*rray by *m*icro-*p*oration), which combines *in vivo* noninvasive, high-throughput time-lapse imaging with excisional biopsies and next generation sequencing to characterize the establishment of the immunological niche and follow its evolution during immunotherapy. STAMP involves the seeding of dozens to hundreds clonal tumors in the superficial dermis of a single mouse ear that can be visualized *in situ* over weeks to months. Using this approach, we found that genetically identical tumors could display surprisingly different immune phenotypes. Although individual tumors of the same array were populated by the same T cell clonotypes, regression or progression of individual tumors were associated with distinct patterns of spatial organization of the T cells. *In situ* imaging of 14K tumors revealed that immune phenotypes were not static over-time but could rather evolve with tumor growth and response to treatment. Therapy-induced or spontaneous early conversion to the immune inflamed phenotype correlated with tumor regression and enhanced cytotoxic T cell activity. Therefore, STAMP provides a flexible approach to study the relationship between tumor evolution, immune cell dynamics, and tumor microenvironment with therapeutic response.

## Main text

The STAMP technique uses an infrared laser (ER:YAG)^9^ to create an array of hundreds of pores in the superficial dermis of a mouse ear pinna (Fig. 1a). Tumor cells from a clonal line (Fig.1a, Extended-Data Fig.1a-c) expressing a fluorescent reporter are seeded into each pore and subsequent tumor growth is monitored over the next 4-6 weeks using live imaging. STAMP can successfully implant orthotopic (melanoma) (Extended-Data Fig.1d-f) or heterotopic (mammary, pancreas, lung and colon carcinoma) tumor cell lines in the dermis of the mouse ear (Fig.1b) or abdomen (Extended-Data Fig.2b). To automatically track large numbers of tumors over time, we developed an image analysis pipeline based on machine learning (Extended-Data Fig.1g-m) that quantifies 1) total tumor burden per animal over time, 2) growth of individual microtumors in the array, and 3) tumor rejection rate.

**Fig 1.**
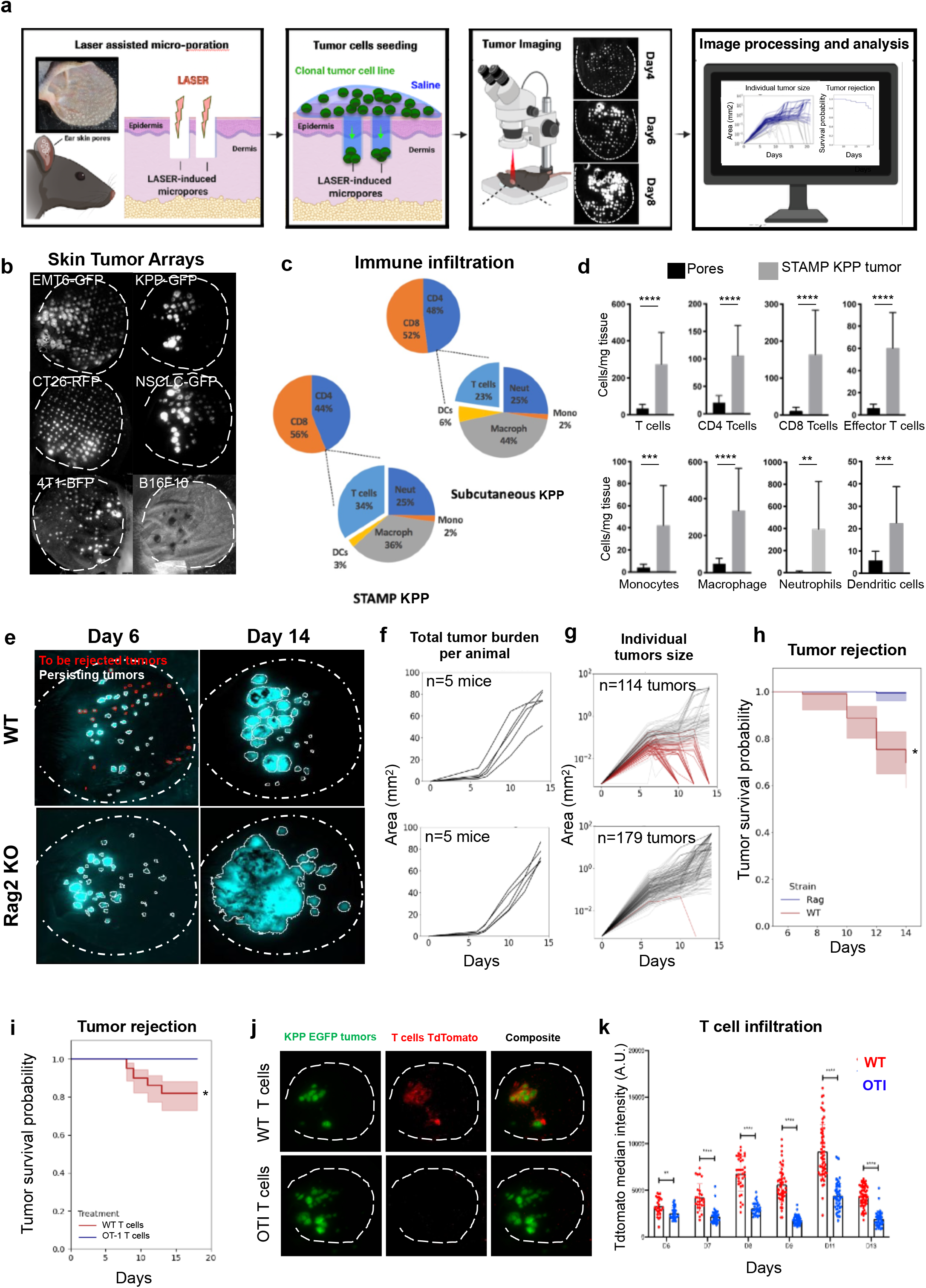
Local T cell-mediated rejection of clonal skin tumor array. **a**, STAMP (<u>*s*</u>kin <u>*t*</u>umor <u>*a*</u>rray by <u>*m*</u>icroporation) workflow. Skin microporation with P.L.E.A.S.E.® Laser Device and subsequent seeding of tumor cell suspension. Individual tumors are longitudinally tracked using epifluorescence microcopy and growth kinetics are analyzed by automated computation. **b**, Representative STAMP tumor arrays of different mouse tumor cell lines at 7 days post tumor implantation for EMT6-GFP, CT26-RFP, 4T1-BFP, KPP-GFP, NSCLC-GFP and 20 days post tumor implantation for B16F10. **c**, Lymphoid and myeloid immune cell profiling of STAMP tumor array and subcutaneous tumors of KPP-EGFP cancer cells. Tumors were harvested18 days post tumor implantation and pooled. n=6 ears, n=3 mice for STAMP. n=5 mice for subcutaneous tumors. **d**, Lymphoid and myeloid immune cell profiling of PBS or KPP-EGFP-seeded micropores. STAMP tumor arrays were harvested 18 days post tumor implantation and pooled. n≧16 ears, n=3 mice. **e**, Representative STAMP tumor arrays of KPP-EGFP in C57Bl6 WT (upper panel) or C57Bl6 Rag2 KO (lower panel) mice at 6 and 14 days post tumor implantation. Red encircled tumors are rejected and white encircled tumors are persistent between time points. n=5 animals. **f**, Automated analysis of growth kinetics of total tumor array area (mm^2^) per animal as described in (e) n=5 animals. **g**, Automated analysis of growth kinetics of individual tumor area (mm^2^) as described in (e-f). n≧114 tumors, 5 animals pooled. Red lines indicate tumors that are rejected, gray lines indicate tumors that persist. **h**, Kaplan Meier survival curves of individual tumors as described in (e-g). n≧114 tumors, 5 animals pooled. (Shaded area=95% confidence interval, log-rank test p-value=2.8×10^-15). **i**, Kaplan Meier survival curves of individual tumors of KPP-EGFP in RAG-2-deficient C57BL/6J mice reconstituted by adoptive transfer of CD3+ T TdTomato+ T cells from either C57BL/6J WT (n≧100) or OT-1+ mice n≧88 tumors. (Shaded area=95% confidence interval, log-rank test p-val=3.1×10^-5). **j**, Representative image of STAMP tumor arrays of KPP-EGFP in RAG-2-deficient C57BL/6J mice reconstituted by adoptive transfer with CD3+ T cells from either C57BL/6J WT (upper panel) or OT-1+ CD4-cre tdTomato+ (lower panel) mice at 9 days post tumor implantation. **k**, T cell infiltration kinetics of individual tumors measured by tdTomato MFI of mice described in (**i**). n≧26 tumors.

### Local T cell-mediated rejection of clonal skin tumor array

To characterize the immune infiltration into STAMP tumors we implanted KPP^10^ clonal cell line (pancreatic ductal adenocarcinoma) subcutaneously or using ear STAMP(Fig.1c) and found similar immune infiltrates regardless of implantation site. Immune cell profiling of ear STAMP tumors was performed by flow cytometry analysis of ears with tumor-bearing vs tumor-free micropores and demonstrated a significant recruitment of myeloid (macrophages, monocytes, neutrophils, dendritic cells) and lymphoid populations (CD4, CD8 T cells) (Fig.1d). To assess the role of adaptive immunity in controlling tumor growth, a STAMP-array of mouse KPP-EGFP tumors was implanted into WT or immunodeficient Rag2-/- mice (Fig.1e). Though WT and Rag-2-deficient animals displayed comparable initial tumor burdens (Fig.1f), after 14 days immunocompetent mice demonstrated a local rejection of 30% of individual tumors from the array, while immunodeficient animals failed to reject tumors (Fig.1g-h).

To assess the role of antigen-specific T cells in this local rejection, we implanted STAMP tumors in Rag-2-deficient mice and simultaneously reconstituted the mice with naïve tdTomato+ T cells from either WT mice, which contain a polyclonal population of T cells, or OT-I mice, which contain a monoclonal population of ovalbumin-specific CD8 T cells (Fig.1i-k). In animals reconstituted with WT tdTomato+ T cells, T cells were recruited to the tumor sites as early as 6 days after tumor implantation and continued to accumulate over time to stimulate local tumor rejection. However, in animals reconstituted with antigen-mismatched tdTomato+ T cells from OT-I mice, T cells were not substantially recruited to tumor sites (Fig.1j-k) and tumors were not rejected (Fig.1i, Extended-Data Fig.1o, Extended-DataVideo1). As KPP-EGFP tumor cells do not express ovalbumin, we can conclude that bystander T cells were not sufficient to promote tumor rejection. Our findings support a role of antigen-specific T cells recruited into STAMP tumors in mediating spontaneous local rejection.

### Heterogeneity and clinical relevance of mouse STAMP tumor-immune phenotypes

To further elucidate drivers of local tumor rejection, we characterized the spatial distribution of T cells in individual tumors. As indicated by fluorescence microscopy, a combination of immune inflamed, excluded and desert tumors were present within each array at all analyzed time points (Fig.2a, Extended-Data Fig.2a-b, Extended-DataVideo2). The ability to monitor individual tumors over time led to the definition of a new late onset phenotype, termed “rejected tumor”, in which EGFP+ tumor cell disappeared leaving a cluster of tdTomato+ T cells for several weeks after tumor rejection. Importantly, by comparison, classical immunophenotyping of tumor biopsies by flow cytometry would not have allowed these diverse phenotypes to have been distinguished (Fig.2c-d, Extended-Data-Fig.2c).

**Fig. 2.**
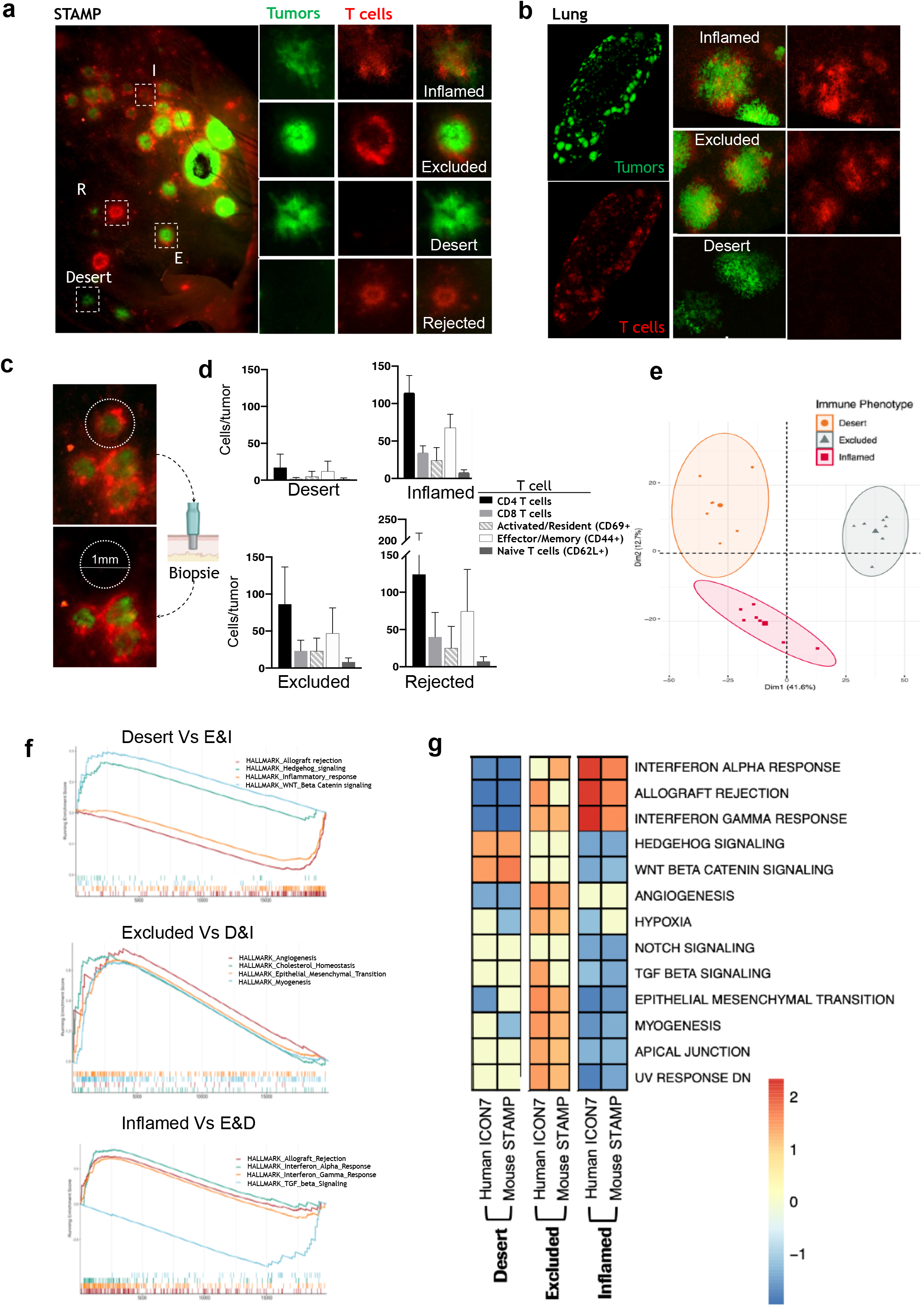
Heterogeneity and clinical relevance of mouse STAMP tumor-Immune phenotypes. **a**, Representative image of STAMP array of KPP-EGFP implanted in C57BL/6J RAG-2-deficient mice at 8 days post tumor implantation, reconstituted by adoptive transfer of CD3+ T cells from C57BL/6J WT CD4-cre tdTomato+ mice (T cells shown in red, KPP-EGFP shown in green). Left panel represents an overview of the entire ear, right panels are enlarged images of individual tumors with diverse immune phenotypes: inflamed, excluded, desert, resolved. n=50 animals. **b**, Representative experimental lung metastases of KPP-EGFP in C57BL/6J RAG-2-deficient mice at 8 days post tumor implantation, reconstituted by adoptive transfer of CD3+ T cells from C57BL/6J WT CD4-cre tdTomato+ mice (T cells shown in red, KPP-EGFP shown in green). Left panel represents an overview of the entire lung lobe, right panels are enlarged images of metastatic foci with diverse immune phenotypes: inflamed, excluded, desert, resolved. n=50 animals. n=5 animals. **c**, Representative STAMP as described in (a), before and after tumor biopsy with 1mm diameter punch biopsy. **d**, Flow cytometry-based T-lymphoid immune cell profiling of biopsied tumors pooled by immune phenotype at day 10. n=6 animals, ≧ 3 pools of tumors. **e**, PCA of bulk RNAseq of individually biopsied tumors with differential immune phenotypes: inflamed, desert, excluded. **f**, Detailed GSEA for selected pathways that are significantly enriched between three different immune phenotypes (inflamed, desert, excluded) from STAMP tumors. **g**, Heat-map comparing the normalized enrichment scores for pathways that are significantly enriched between the three different immune phenotypes (inflamed, desert, excluded) from either the human ICON7 clinical trial or the mouse STAMP tumors.

As the tumors from the array share the same clonal origin and are implanted simultaneously at a single site in a single mouse, it was remarkable that distinct tumor immune phenotypes could develop. We therefore next confirmed that heterogeneous T cell distribution in tumor arrays was not an artifact of the STAMP method. For this purpose, we examined if different immune phenotypes would also develop using a model dependent on spontaneous tumor seeding. The immune phenotypes of lung metastatic tumors were monitored following intravenous seeding of clonal KPP-EGFP tumor cells in Rag2-/- mice reconstituted with TdTomato+ T cells. As observed in STAMP, the full spectrum of tumor-immune phenotypes was observed in lung nodules imaged after optical clearing (Fig.2b, Extended-DataVideo3). Thus, our findings demonstrate clonally-derived tumor cells have the unanticipated ability to develop heterogeneous immune phenotypes independently of variations in either host or tumor genetics

To elucidate the basis for the diversity of immune phenotypes, we next explored the biological pathways associated with each tumor niche. STAMP arrays were first imaged to classify tumors by immune phenotype and individual tumors were then isolated by micro-punch biopsy for bulk RNA sequencing (Fig.2c-e). Principal Component Analysis revealed that different immune phenotypes constitute distinct transcriptional clusters (Fig.2e). Differential expression analysis of genes (Extended-Data Fig2.d) and pathways (Fig.2f) identified gene signatures enriched in immune inflamed, excluded, and desert tumors that were stable over time (Extended-Data Fig2.e). The gene signatures enriched for each immune-phenotype of STAMP tumors were then compared with pathways enriched for tumor-immune phenotypes in clinical samples using bulk RNAseq of human ovarian (Fig.2g) and bladder FFPE tumors from clinical trials (ICON7 phase III and IMvigor210 phase II, respectively) (Extended-Data Fig.2e-f) that were classified as desert, excluded, or inflamed based on CD8 and pan-cytokeratin staining.

Inflamed tumors from both mouse STAMP and human tumors were enriched for IFNα, IFNγ, and allograft rejection signatures and were also characterized by a downregulation of the TGF-β^17^ and NOTCH pathways. As previously described, T cell and interferon signatures are both well validated drivers of the immune inflamed phenotype^18^ as is downregulation of TGF-β pathway^11^. Immune excluded tumors were enriched for epithelial-mesenchymal transition (EMT), myogenesis, angiogenesis, hypoxia and UV response signatures. EMT, myogenesis and angiogenesis enrichment in excluded tumors is consistent with the enrichment in stromal and myofibroblast components described previously^12,13^. Finally, immune desert tumors were enriched for WNT/β-catenin^22^ and Hedgehog pathways^23^ and showed a downregulation of immune signatures including allograft rejection, T cell, and interferons. WNT/β-catenin has previously been associated with poor immune cell infiltration^14,15^. The overlap of significant pathways enriched in mouse STAMP tumors and human tumors of the same immune phenotypes held across different human cancer types (ovarian, bladder and lung) (Extended-Data Fig.2e-f and data not shown) Thus, the similarity of gene signatures associated with the three immune phenotypes in both human and STAMP tumors suggests that STAMP may recapitulate the general mechanisms underlying this aspect of tumor diversity observed clinically.

### Immune phenotype determines T cell function regardless of T cell clonotype

Given the heterogeneity in spatial organization of T cell infiltrates among individual tumors, we next asked if these variations correlated with qualitative differences in local T cell responses. Although each tumor was derived from a common cell line and the T cells likely derived from the same draining lymph nodes, it was possible that differences in T cell infiltration and tumor growth control reflected differences in local T cell receptor clonotype profiles. To this end, we performed 5’single cell RNAseq and TCR-seq of tumor punch biopsies harvested from the same mouse 8 days after tumor implantation; individual tumors of the same immune phenotype were pooled to provide sufficient numbers of cells for the analysis.

Single cell analysis identified twelve clusters of T cells which included four CD8 T cell clusters: two CD8 T effector cell clusters, one CD8 T_RM_ cell cluster and one mitotic CD8 T cell cluster. The remaining eight sub-clusters were CD4 T cell clusters, including two mitotic CD4 T cell clusters, one Treg cluster and five CD4 effector T cells clusters (Fig.3a,b; Extended-Data Fig.3a-d). Interestingly, we did not observe any difference in absolute or relative abundance of T cell subsets across tumors with inflamed, excluded and resolved phenotypes (Fig.3c,d; Extended-Data Fig.3e,f). Desert tumors, as expected, were devoid of most T cells populations.

The relative and total abundance of T cell clonotypes was also shared across inflamed, excluded and rejected phenotypes (Extended-Data Fig.3g-j). We identified seven immunodominant clonotypes each of them present at comparable frequencies among inflamed, excluded, and rejected tumors (Fig.3d,e; Extended-Data Fig.3h-i). However, we found that identical T cell clonotypes exhibit divergent gene expression profiles when present in tumors with differing immune phenotype. Differential gene and pathway analysis of immunodominant T cell clonotypes showed that T cells clones present in inflamed tumors were characterized by signatures indicating an increased translation and mitochondrial biogenesis compared to the same T cells clones located in excluded tumors. (Fig.3f; Extended-Data Fig3k,n).

Because mitochondrial dynamics control T cell fate^16–18^ and translational activity correlates with the activation and differentiation state of the effector T cells^19^, we investigated the functionality of the T cells present in immune inflamed and immune excluded tumors. We developed a clonal KPP tumor cell line expressing the calcium sensor^20^ GCaMP6 that emits green fluorescence when the intracellular Ca^2+^ concentration rises, as occurs upon effector T cell attack and membrane perforation. Repetitive T cell attacks highlighted by repetitive Ca^2+^ flashes in tumor cells were followed up by tumor cell death highlighted by Propidium Iodide uptake (Fig.3g-i; Extended-Data Fig.3o-s). Therefore, the occurrence of green flashes in tumor cells correlate with T cell cytolytic activity. We validated the sensitivity and the specificity of calcium flashes in response to T cell killing *in vitro* using KPP organoids (Fig.3h,i; Extended-Data Fig.3p-s; Extended-DataVideos4,5) and *in vivo* using STAMP microtumors (Extended-Data Fig.3t,u; Extended-DataVideos6-8). We then quantified the difference in Ca^2+^ flashes between immune inflamed and immune excluded tumors from STAMP arrays at 8 days after implantation and further correlated it with their tumor growth rates (Fig.3j-n; Extended-DataVideos9-10). The flashing index of inflamed tumors was found to be significantly higher as compared to excluded tumors (Fig.3m) despite comparable total T cell abundances (Fig.3l). Indeed, a high flashing index corresponded to an immune inflamed phenotype and a slow tumor growth rate (Fig. 3n; Extended-Data Fig.3v,w).

**Fig. 3.**
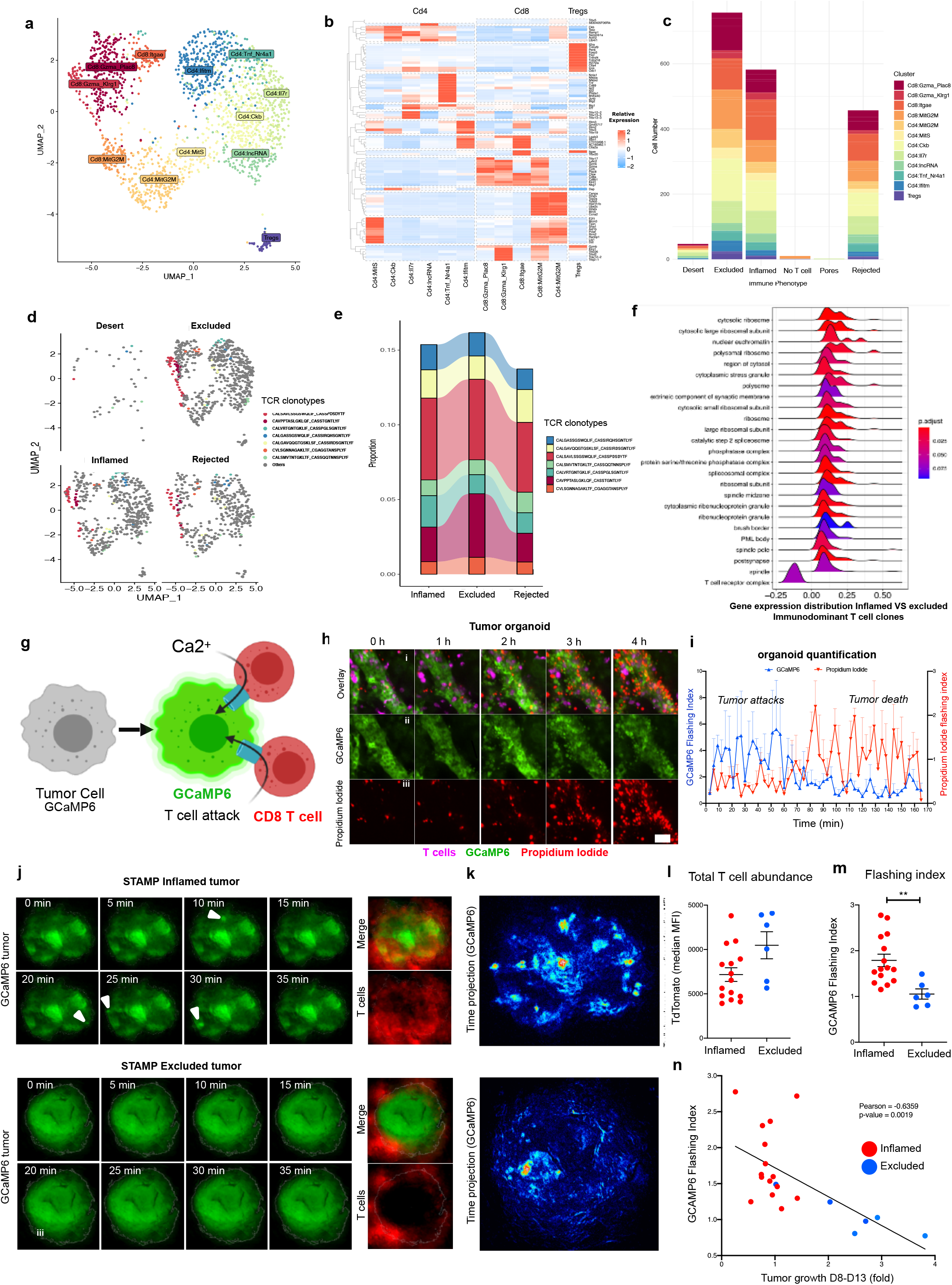
Immune phenotype determines T cell function regardless of T cell clonotype. **a**, UMAP visualization of tdTomato+ T cell clusters from pooled STAMP tumor biopsies. **b**, Heatmap of the average normalized expression of top 10 positive marker genes for each T cell cluster. **c**, Cell numbers of each T cell sub-clusters per tumor immune phenotype. **d**. UMAP visualization of the T cell clusters per tumor immune phenotypes with overlay of the top seven immunodominant clonotypes **e**, Relative abundance of the top seven immunodominant clonotypes per Immune phenotypes. **f**, Expression distribution of the core enriched genes for the top 25 enriched pathways comparing immunodominant T cell clones present in Inflamed VS excluded tumors, corrected for their T cell subcluster identity **g**, cytotoxic T cells creating Ca2+ permeable perforin pores in the tumor cell membrane leading to tumor cell killing. Ca2+ influx activates green fluorescence of GCaMP6 calcium sensor. **h**, Representative in vitro time-lapse images of GCaMP6-expressing tumor organoids under T cell attack. (i) Composite of GCaMP6 (green), propidium iodide (red) and T cell fluorescence (magenta) (ii) single channel of GCaMP6 fluorescence, (iii) single channel of propidium iodide fluorescence, n=3. **i**, quantification of calcium flashes and propidium iodide accumulation during T cell-mediated tumor cell killing of organoids. n=3. **j**, Left panel**-**Representative *in vivo* time-lapse images of GCaMP6-expressing inflamed (upper panel) or excluded (lower panel) STAMP tumors, as described in (g). Right panel-composite of tumor and T cell fluorescence to highlight tumor Immune phenotype. GCaMP6 tumor: green and tdTomato T cell: red **k**, Time projection of maximum minus mean GCaMP6 fluorescence of the tumor described in (j) **l**, Total T cell abundance (including adjacent area) of individual inflamed and excluded tumors from GCaAMP6 expressing KPP-STAMP arrays n≧6. * p-value ≦ 0.05, unpaired two-sided T-test. **m**, Ca2+ influx indices of inflamed and excluded tumors described in (l). n≧6. ** p-value ≦ 0.01, unpaired two-sided T-test. **n**, Correlation analysis of flashing indices and tumor growth fold change (Day8-to-Day13) of inflamed and excluded tumors described in (l).

Taken together, these results indicate that T cells of the same TCR clonotype exhibit different functional capacity when localized to immune inflamed vs immune excluded tumors, emphasizing a determinative role for the tumor microenvironment in shaping the activity and fate of endogenous effector T cells.

### Early transition to the inflamed phenotype predicts tumor response to immunotherapy

Finally, we explored how the spatial distribution of T cells predict tumor progression or rejection during Immunotherapy. Although human and mouse tumors are well known to adopt one or another immune phenotype, it remains unclear whether these states are stable or dynamic. STAMP provides an opportunity to determine how immune phenotypes evolve with immunomodulatory therapy. As immune excluded and immune desert STAMP tumors exhibited an upregulation of the TGFb pathway (Fig.2g), we hypothesized that TGFb could restricts T cell entry into the tumor parenchyma. It has been previously shown that both T cell recruitment and therapeutic benefit can be elicited in mice by treatment with TGFb antagonists in combination with checkpoint blockade anti-PD-L1^21 11^. We treated STAMP tumor-bearing mice with anti-PD-L1, anti-TGF-β or a combination of both. While treatment with anti-PD-L1 or anti-TGF-β alone failed to improve tumor rejection rates, combination therapy led to a 73.3% overall response rate (ORR) in immunocompetent mice (Extended-Data Fig.4a-c) and 69% O.R.R in RAG2-/- mice reconstituted with adoptively transferred T cells (Fig.4a-b, Extended-Data Fig.4d).

As the combination therapy was able to reject tumors that were larger than those rejected spontaneously in the control arm (Fig.4b-c), we next quantified T cell abundance in tumors as a function of time. We performed daily imaging of STAMP KPP-EGFP tumors implanted in RAG2-/- mice reconstituted with tdTomato+ T cells and treated with anti-TGFβ and anti-PD-L1. As shown in Fig. 4d, tumors in combination-treated mice exhibited a significant increase in total T cells starting at Day 10. Although STAMP tumors in mice treated with anti-PD-L1 alone also exhibited increases in CD8 T cell infiltration (Extended-Data Fig.4e), only the combination therapy led to tumor rejection (Fig.4a).

**Fig. 4.**
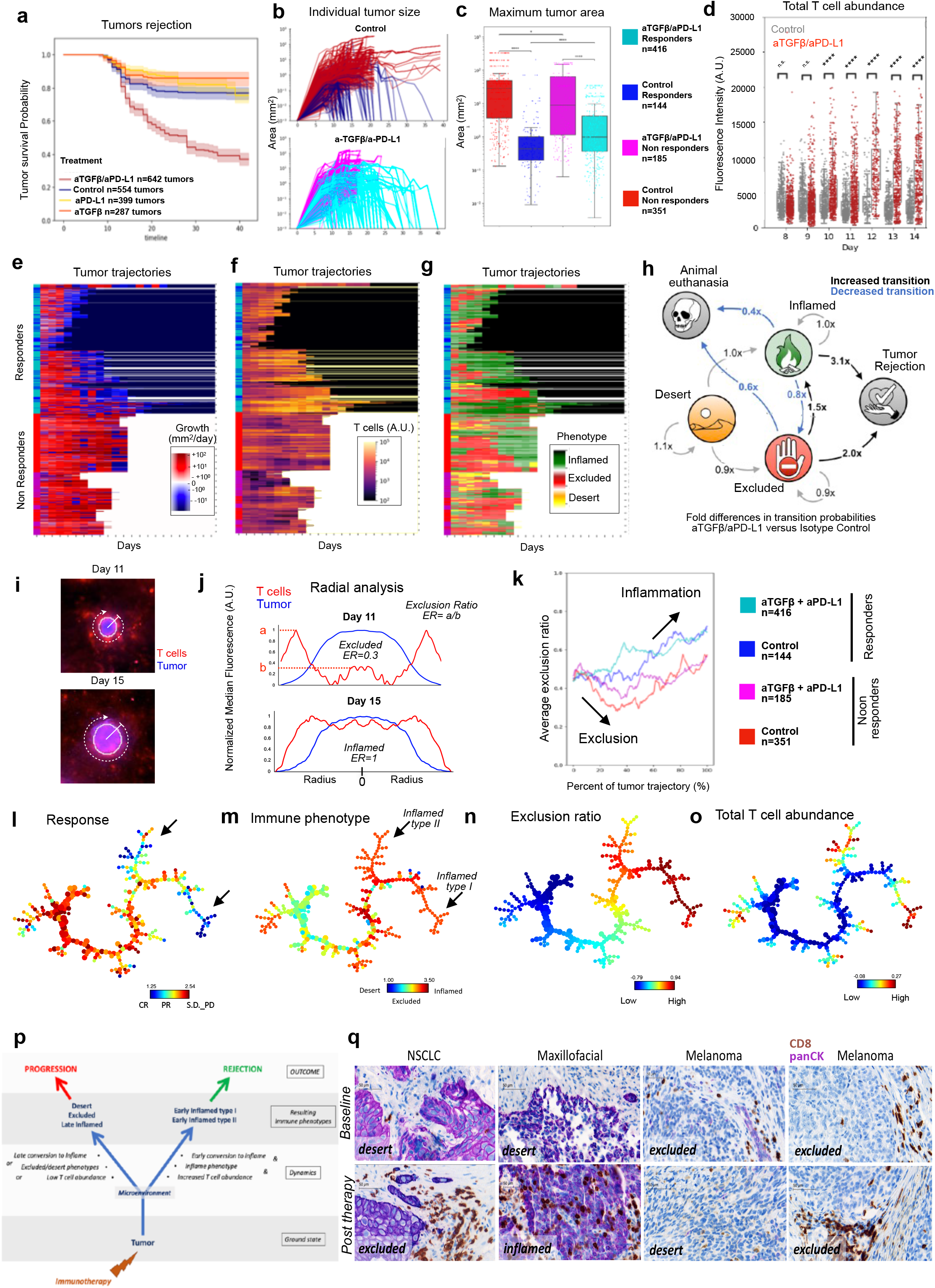
Early transition to an inflamed phenotype predicts response to Immunotherapy. **a**, I Kaplan Meier survival curves of individual tumors implanted in Rag2 KO animals reconstituted with tdTomato T cells and and treated at day 1 post-implantation with isotype control antibodies, anti–TGF-β, anti–PD-L1, or a combination of anti–PD-L1 with anti–TGF-β. n≧287 tumors per group and 10-12 animals per group. Shaded area=95% confidence interval, log-rank test p-value n.s. >0.5, ****<0.0001. **b**, Individual tumor growth kinetics (mm^2) of control and combination anti–PD-L1 with anti–TGF-β treated tumors. n≧287 tumors per group and 10-12 animals per group. **c**, Maximum area achieved of Responders (C.R + P.R) and Non Responders (S.D+P.D) tumors for combination anti–PD-L1 with anti–TGF-β versus isotype control treated animals as described in (b). **d**, Total T cell abundance over time (tdTomato MFI per tumor) for individual tumors for isotype control treated tumors (gray dots showed as reference) and combo anti–PD-L1/ anti–TGF-β treated tumors (red dots). **e-g**, Hierarchical clustering of individual tumor immune trajectories for isotype control treated tumors and combo anti–PD-L1/anti–TGF-β treated tumors. black=tumor resolved, white=mouse euthanized. Combo Responders (C.R. + P.R.) in cyan, combo Non Responders (S.D. + P.D.) in magenta. Control Responders (C.R. + P.R.) in blue, control Non Responders (S.D. + P.D.) in red. **e**, Tumor growth rate (mm2/day) over time **f**, T cell abundance (TdTomato MFI per tumor over time) **g**, Tumor Immune phenotype over time **h**. Markov chain showing the fold difference between transition matrices for isotype control versus combination anti–PD-L1 with anti– TGF-β treated animals as described in (b), **i**, Epifluorescence images of the same individual tumor at day 11 and day 15 with overlaid of radial analysis. Tumor was selected from an isotype control treated animal from the experiment described in (a). Tumor=blue, T cells=red. **j**, quantification of median radial fluorescence profile for individual tumor shown in (i) at day 11 (top) and day 15 (bottom) to distinguish T cell excluded (low T cell exclusion ratio) and inflamed phenotype (high T cell exclusion ratio) **k**, average exclusion ratio over normalized tumor trajectory for Combo Responders (C.R. + P.R.) in cyan, combo Non Responders (S.D. + P.D.) in magenta. Control Responders (C.R. + P.R.) in blue, control Non Responders (S.D. + P.D.) in red. **l**, SPADE tree representing the layout of 11K individual tumors across days 6-17 for all treatment groups, in the space of the four parameters (Total T cell abundance, T cell abundance in the tumor core; tumor exclusion ratio; Tumor size). SPADE trees were color-coded according to the response to therapy (panel **l**), Immune phenotype (panel **m**) exclusion ratio (panel **n**) and T cell abundance (panel **o**). Categorical variables such, e.g. phenotype or response, were converted to numerical variables that were further used to color the SPADE tree. Size of the dots is proportional to the number of tumors **p**, graphical summary of Immune dynamics leading to outcome **q**, Histology of patient paired tumor biopsies at baseline and at progression after treatment with checkpoint blockades. CD8 staining in brown, pan-Cytokeratin staining in magenta.

Having shown that a simple increase in total T cells at the tumor site was insufficient to induce efficient tumor rejection, we next sought to identify other immunological features that precede efficient rejection. We therefore examined the spatial distribution of T cells over time in response to the combination therapy using a high content image analysis pipeline to automatically classify tumors as immune inflamed, excluded or desert. T cell phenotype classifications could then be overlaid with tumor growth curves to create a trajectory representation of tumor fate (Extended-Data Fig.4f-h). Tumor trajectories were plotted using hierarchically clustered heatmaps illustrating tumor growth rate (Fig.4e), median T cell abundance (Fig.4f), and tumor immune phenotype over time (Fig.4g). The analysis of 1.2K individual tumors implanted in 20 animals showed that responding tumors (C.R.+P.R.) in both combination therapy-induced rejections (turquoise) and spontaneous rejections (dark blue) shared similar trajectories over time, both showing a negative growth rate and higher degree of T cell recruitment prior to tumor rejection.

To further characterize dynamic changes between immune phenotypes, we developed a Markov chain model to compute changes in transition probability among the different states: desert, excluded, inflamed, rejected tumor and animal euthanasia (Fig.4h, Extended-Data Fig.4i-k, Extended-DataTable1). We observed that responding tumors (CR+PR) were in general more likely to transition from desert to inflamed and from excluded to inflamed than non-responding tumors (SD+PD) regardless of the treatment (Extended-Data Fig.4i,j). The combination treatment relative to control increased the probability of transition from excluded to inflamed and decreased the probability of reverse transition from inflamed to excluded (Fig.4h). Additionally, the probability of an inflamed or an excluded tumor becoming resolved was increased relative to control if treated with combination immunotherapy (Fig.4h). These results suggest that differences in tumor outcome may be associated with an increased probability of a tumor becoming inflamed, and that such transitions may be promoted by combination immunotherapy

To better understand which portion of the tumor trajectory determines tumor outcome, we used a radial analysis to calculate an exclusion score (maximum T cell abundance in the tumor core/maximum T cell abundance at the tumor periphery) for tumors over time (Fig.4i-k, Extended-Data Fig.4l). Although tumors increased their inflammatory features over time, the first 20% of the trajectory of non-responding tumors (SD+PD; red, magenta) became substantially more excluded than responding tumors (CR+ PR; turquoise, blue) (Fig 4.k). Thus, if an individual tumor became more excluded early in its trajectory, that tumor would be less likely to be rejected. This analysis highlights the importance of early T cell infiltration into the tumor core, as a transition to an inflamed phenotype late in a tumor’s lifetime is less efficient at inducing tumor regression (Fig.4k).

Finally, to identify deeper patterns in the dataset, we performed multivariate clustering analysis using UMAP (Extended-Data Fig.4m-o) and SPADE tree (Fig.4l-o, Extended-Data Fig4r-v) of 11K images of individual tumors acquired between Day 6-17 (Extended-DataTable2) to observe treatment-specific immunomodulation over time (Extended-Data Fig.4n,o) and to further quantify it using the Earth Moving Distance analysis (dissimilarity score) for each treatment group (Extended-Data Fig.4o-q). This multivariate analysis revealed that T cell dynamics of the anti-PD-L1/anti-TGFb combo treatment group has significantly smaller dissimilarity to the Immune dynamics of responder tumors than any of the other treatment groups over time. Additionally, SPADE trees generated a hierarchical clustering that revealed patterns of heterogeneity within the treatment groups (Fig.4l-o, Extended-Data Fig.4r,v) previously ignored by a simple phenotype classification (inflamed, excluded, desert). As shown in Fig.4l the responders group splits into two subpopulations of inflamed tumors (arrows): type I, which has a higher exclusion ratio and total T cell abundance, and type II, which had lower values (Fig.4m-o). Both subtypes had high probability of response to therapy.

As summarized in Fig.4p, STAMP reveals a spatial and temporal control of Immune-mediated tumor rejection relying on 1) increased total T cell abundance at the tumor site, 2) transition to an inflamed phenotype, 3) earliest T cell infiltration in the tumor core. STAMP could also be used to identify heterogenous patterns of response to therapeutics facilitating future exploration of novel classes of tumors.

### Evolution of tumor immune phenotype in human cancers

The analysis of STAMP tumors indicated that tumor immune phenotypes were capable of dynamic evolution that determine their response to therapy. To assess whether a similar situation might exist for human tumors, we analyzed on treatment biopsies taken from six patients with various solid tumor indications both before and during treatment with checkpoint inhibitors (Extended-DataTable3). The distribution of CD8 T cells was determined by immunohistochemistry relative to the tumor nest. As shown in Fig.4q, it is apparent that the immune phenotype of a tumor can convert over time with therapeutic intervention, and not always from less to more inflamed. In the paired samples shown, examples of transitions immune desert to excluded (NSCLC), desert to inflamed (maxillofacial), excluded to desert (melanoma I) and excluded to excluded (melanoma II) can be seen. Understanding the spontaneous or CIT-induced evolution of tumor Immune phenotypes in the clinic will be an important consideration in predicting the likelihood of clinical response. Further modeling of tumor-immune dynamics by STAMP and integration of multiple spatial and temporal variables^22^ may provide insight into the mechanisms determining tumor fate and provide the biomarkers that may guide therapy in human cancer patients.

## Methods

### STAMP (skin <u>*t*</u>umor <u>*a*</u>rray by <u>*m*</u>icroporation) implantation

Experimental procedures were performed in 8-12-week-old male mice under anesthesia (60-100 mg/kg ketamine & xylazine 5-10 mg/kg, intraperitoneal injection). Prior to microporation, the ear hair was removed with Veet® depilatory cream then rinsed with water, dried, and the dorsal side of the ear was immobilized with double-sided tape to expose the ventral side. Microporation at 71 µm depth and 9% pore density is performed by applying the P.L.E.A.S.E laser device (Pantec Biosolutions, Liechtenstein) to the ventral side of the ear using a custom program previously validated for the cell line.

After the microporation process, 150 µL of a tumor cell suspension (EMT6-EGFP, CT26-RFP, 4T1-mTagBFP2, KPP-EGFP, KPP-mTagBFP2, NSCLC-EGFP or B16F10) at 40-80×10^6^cells/mL diluted in phosphate buffered saline was applied to the ear covering the ventral side. Cells were incubated for 30 min then the excess cell suspension was removed, pores were covered with Matrigel (Corning, NY, USA) and incubated for 15 min until matrigel polymerized. Microtumor growth becomes evident 5 to 12 days post tumor implantation, depending on the cell line.

### Mice

The Genentech institutional animal care and use committee responsible for ethical compliance approved all animal protocols. Mice were housed under specific pathogen-free conditions at Genentech animal facility, South San Francisco, CA. Animals between 8–12 weeks old that appeared healthy and free of obvious abnormalities were used for the study.

B6.Cg-Foxn1nu/J (Stock No. 000819), C57BL/6-Tg(CAG-EGFP)1Osb/J (Stock No. 003291), and C57BL/6J (Stock No. 000664) animals were purchased from the Jackson Laboratories (ME, USA). B6.129S6-Rag2^tm1Fwa^ N12 (Model No. RAGN12), C.Cg/AnNTac-Foxn1^nu^ NE9 (Model No. BALBNU-M), and BALB/cAnNTac (Model No. BALB-M) animals were purchased from Taconic Biosciences (CT, USA).

CD4.cre.tg Rosa26.LSL.tdTomato.cki OT-I.TCR.tg (OT1-/-and OT1+/+) animals and E8I.CD8A.IRES.GFP.Cre.tg Rosa26.LSL.tdTomato.cki animals were bred in house.

### Subcutaneous tumors

For subcutaneous tumor inoculation, mice were injected subcutaneously with 0.5×10^6^ KPP-EGFP cells in 100 μL of a 1:1 dilution of phosphate buffered saline and Matrigel (Corning, NY, USA).

### CIT treatment

Animals were implanted with STAMP microtumors as described previously. Mice were distributed into treatment groups to exclude cage effects and when possible, account for differential initial tumor growth. Treatment was initiated one day post tumor implantation for immunodeficient mice or 7 days post tumor implantation for immunocompetent mice. Animals were treated every other day with isotype control antibodies (anti-gp120, mouse IgG clone 3E5, 10 mg/kg), anti–PD-L1 (mouse IgG1 clone 6E11, 10 mg/kg first dose followed by 5 mg/kg thereafter), anti–TGF-β (mouse IgG1 clone 1D11, 10 mg/kg), or a combination of anti–PD-L1 with anti–TGF-β. 5 days after treatment initiation, mice were imaged daily on a M205FA stereoscope with a 1.0x PlanApo lens (10450028; Leica Microsystems, Germany) and ORCAII Digital CCD (Hamamatsu Photonics, Japan) to monitor tumor growth and T cell progression. For selected experiments, immunodeficient mice were injected intravenously with 4×10^6^ isolated T cells-Tomato from CD4.cre.tg_Rosa26.LSL.tdTomato.cki_OT-I.TCR.tg as described below.

### Metastasis mouse model and imaging of whole mount tissue

RAG-2-deficient mice were injected intravenously with 0.1×10^6^ KPP-EGFP cells and 4×10^6^ tdTomato+ CD3+ T cells isolated as described below. 8 days post intravenous injection, mice were euthanized, and 20 mL of cold PBS/heparin 5U/ml solution was perfused directly into the right ventricle using a 27-gauge needle. Lungs were isolated by dissection and tissues were fixed using 4% paraformaldehyde in phosphate buffered saline^23^. Tissue clearing was performed using the FluoClearBABB approach^24^ and whole mount images were then acquired using a SP8 microscope equipped with a white light laser and a HCX APO L 20x/0.95 IMM lens (Leica Microsystems, Germany). Imaging data was analyzed on a workstation (Thinkmate, MA, USA) using Imaris software (Bitplane, United Kingdom).

### T cell isolation

T cells were isolated from C57BL/6J or CD4.cre.tg Rosa26.LSL.tdTomato.cki OT-I.TCR.tg (OT1-/-and OT1+/+) mice. Spleens were collected and dissociated with the end of a plunger from a 1 mL syringe into 10 mL of PBS before filtration through a 70 μm cell strainer. T cells were isolated for intravenous injection using an EasySep™ Mouse T Cell Isolation Kit (Stemcell Technologies, Canada). 4×10^6^ isolated T cells were injected intravenously per mouse.

To generate CTLs from OT-I mice, splenocytes were isolated and stimulated with 10 nM OVA257-264 peptide (AnaSpec, CA, USA) in complete media containing Gibco RPMI 1640 (Thermo Fisher Scientific, MA, USA) with 10% Gibco fetal calf serum (Thermo Fisher Scientific, MA, USA), 2 mM L-glutamine, 50 U/mL Gibco penicillin/streptomycin (Thermo Fisher Scientific, MA, USA), and 50 μM β-mercaptoethanol. Following 3 days of stimulation, cells were resuspended in complete media with 10 IU/mL recombinant human IL-2 (rHIL-2). CTLs were kept at a density of 0.5×10^6^ cells/mL and fresh complete media with rHIL-2 was added every 48 hours. CTLs were used between 6 and 8 days after primary *in vitro* stimulation.

### Flow cytometry

Ear tissue was isolated using a 1 mm Miltex sterile disposable biopsy punch with plunger (Integra Biosciences, NH, USA) from animals bearing STAMP microtumors or mock-implanted control mice at either 8 or 18 days post tumor implantation. Tumors were digested in 500 µL (STAMP microtumors) or 3000 mL (subcutaneous) of phosphate buffered saline containing 0.1 mg/mL DNAse I (Roche, Switzerland) and collagenase D at 1 mg/mL (Roche, Switzerland) for 30 min at 37°C to obtain a single cell suspension.

For surface staining, cells from digested tumors were incubated with Fc block (5 µg/mL; BD Biosciences, CA, USA; clone 2.4G2) and stained with antibody mix for 30 min and Viability Dye eFluor™ 780 (eBioscience, CA, USA).

Antibodies were used at 1:200 dilution. Anti-mouse CD19 (clone B4), anti-mouse I-A/I-E (clone M5/114.15.2), anti-mouse F4/80 (clone BM8), anti-mouse/human CD11b (clone M1/70), anti-mouse Ly-6G (clone 1A8), anti-mouse Ly-6C (clone HK1.4), anti-mouse CD69 (clone H1.2F3), anti-mouse CD25 (clone PC61), anti-mouse CD4 (clone RM4-5), anti-Mouse CD62L (clone MEL-14), and anti-mouse/human CD44 (clone IM7) antibodies were purchased from Biolegend (CA, USA). CD45 anti-mouse (clone 30:F11), anti-mouse CD86 (clone GL1), CD11c anti-mouse (clone N418), and anti-mouse CD8a (clone 53-6.7) antibodies were purchased from ThermoFisher Scientific (MA, USA). CD3 anti-mouse (clone 17A2) antibody was purchased from BD Biosciences (CA, USA).

Live singlets cells subsets CD45+ were gated as follow: MHC class II+ CD11c+ F4/80-as DC or CD103+ or CD86+ activated DC; CD11c-CD11b+ F4/80+ as macrophages, CD11b+ Ly6G+ Ly6C^int^ as neutrophils; CD11b+ Ly6G^low^ Ly6C+ as monocytes; CD3+ T cells were divided as CD3+ CD4+ T cells, CD3+ CD8+ T cells, CD3+ CD69+ activated/resident T cells, CD3+ CD44+ CD62L-effector/effector memory T cells or CD3+ CD44-CD62L+ naive T cells.

Flow cytometry data were collected with a BD LSRFortessa cell analyzer (BD Biosciences, CA, USA) and analyzed using FlowJo Software (Version 10.2; FlowJo LLC, OR, USA).

### Whole Exome Sequencing Analyses

We conducted whole exome sequencing of DNA isolated from the KPP-EGFP cell line harvested at the end of a culture passage, under the conditions for the experiments of this work, in order to ascertain the absence of major sub-clones explaining the diversity of immune phenotypes observed. A spleen sample from the mice strain used in our experiment were used as matched normal. Exome capture libraries were sequenced on HiSeq 2500 (Illumina) to generate 2 × 75 bp paired-end data, from the sequenced reads, variants were called using the following workflow. Sequencing reads were mapped to UCSC mouse genome (GRCm38) using BWA software^25^ set to default parameters. Local realignment, duplicate marking, and raw variant calling were performed according to GATK best practices^26^. Somatic variant calling on tumor and its matched normal BAM file was performed using Strelka^27^. The resulting VAFs allowed us to construct histograms to demonstrate clonality. Additionally, the R package neutralitytestr (https://github.com/marcjwilliams1/neutralitytestr) was used to show that low frequency variants detected are indeed part of the neutral drift that any cultivation process entails.

### STAMP microtumor sample preparation for bulk RNAseq of tumor biopsies

FOXN1-deficient nude mice with adoptively transferred tdTomato+ CD3+ T cells bearing KPP-EGFP STAMP microtumors were biopsied 8 days post tumor implantation using a 1 mm Miltex sterile disposable biopsy punch with plunger (Integra Biosciences, NH, USA). Each tissue biopsy was transferred to a separate 1.5 mL tube (Eppendorf, Germany) containing 0.25 mL Invitrogen TRIzol Reagent (Thermo Fisher Scientific, MA, USA). Biopsies were incubated in TRIzol for 5 min with intermittent vortexing. 50 µL chloroform (MilliporeSigma, MA, USA) was added to the homogenate, vortexed for 20 sec, and incubated at 20-25°C for 2-3 min. To accelerate phase separation, samples were centrifuged at 10,000xg for 18 min at 4°C. The aqueous (top) phase was removed by aspiration and transferred to a clean 1.5 mL tube (Eppendorf, Germany). A volume of 100% RNAse-free ethanol (MilliporeSigma, MA, USA) equal to the volume of the aqueous layer was added, and the RNA was further isolated using a RNeasy Micro Kit (Qiagen, Germany). Alternatively, an individual tumor biopsy was immersed in RNA later and the RNA further extracted using a RNeasy Mini Kit (Qiagen, Germany).

### STAMP microtumor bulk RNAseq analysis

RNA-sequencing data were analyzed using HTSeqGenie pipeline in BioConductor^28^ as follows: first, reads with low nucleotide qualities (70% of bases with quality ¡23) or matches to rRNA and adapter sequences were removed. The remaining reads were aligned to the mouse reference genome GRCm38.p5 using GSNAP (PMID:20147302, 27008021) version ‘2013-10-10-v2’, allowing maximum of two mismatches per 75 base sequence (parameters: ‘-M 2 -n 10 -B 2 -i 1 -N 1 -w 200000 -E 1 –pairmax-rna=200000 –clip-overlap’). Transcript annotation was based on the Gencode genes database^29^. To quantify gene expression levels, the number of reads mapping unambiguously to the exons of each gene was calculated. The resulting count matrix was filtered for lowly expressed genes keeping the genes with at least 0.2 counts per million in more than six samples. Principal component analysis was performed with the factoextra R package. Differential gene expression analysis was performed using the voom and limma R packages^30^. To detect signature differences amongst immune phenotypes, our contrasts compared the immune phenotype at hand with the average of the other two. Volcano plots were constructed using the EnhancedVolcano R package (https://github.com/kevinblighe/EnhancedVolcano). Gene set enrichment analysis was performed on the log Fold Changes with using the GSEA function with ClusterProfiler R package the Hallmark Gene Set Collection^31^. Pathways were considered significant if their multiple testing corrected p-value was less that 0.2. The same analysis was performed on the clinical trial RNA seq data. Heatmaps of the normalized enrichment score were constructed for those significant pathways was constructed using the ComplexHeatmap R package^32^.

### Sample preparation of sorted STAMP microtumors for scRNAseq

FOXN1-deficient nude mice with adoptively transferred tdTomato+ CD3+ T cells bearing KPP-EGFP STAMP microtumors were biopsied 8 days post tumor implantation using a 1 mm Miltex sterile disposable biopsy punch with plunger (Integra Biosciences, NH, USA). 3-6 pooled tissue biopsies were moved into a precooled 1.5 mL tube (Eppendorf, Germany) containing 300µL digestion cocktail consisting of Gibco RPMI 1640 (Thermo Fisher Scientific, MA, USA), 0.1% Gibco fetal calf serum (Thermo Fisher Scientific, MA, USA), 0.1mg/ml Liberase TM (Roche, Switzerland), 0.1mg/ml DNAse I (Roche, Switzerland), 32uM Gibco Actinomycin D (Thermo Fisher Scientific, MA, USA). Tissues were incubated for 30 min at 37°C and 950x RPM on a Thermoblock (Eppendorf, Germany) and mechanically dissociated every 10 min with a pipette. To quench the digestion, the cell suspension was filtered through a 40 µm mesh into pre-cooled FACS filter tube containing quenching buffer of Gibco fetal calf serum (Thermo Fisher Scientific, MA, USA) with 32 uM Gibco Actinomycin D (Thermo Fisher Scientific, MA, USA). The cell suspension was centrifuged at 350xg for 8 min and resuspended in 400 µL Gibco RPMI 1640 (Thermo Fisher Scientific, MA, USA) with 5 µM CalceinBlue (Invitrogen, CA, USA) and a 1:200 dilution of Molecular Probes Fixable Live/Dead Near-IR Dead Cell Stain Kit (Thermo Fisher Scientific, MA, USA). FACS was performed to isolate Calcein Blue-positive and L/D NearIR-negative cells into a precooled 1.5ml Eppendorf tube containing 750µL collection buffer consisting of Gibco RPMI 1640 (Thermo Fisher Scientific, MA, USA), 10% Gibco fetal calf serum (Thermo Fisher Scientific, MA, USA), 32 uM Gibco Actinomycin D (Thermo Fisher Scientific, MA, USA). The cell number and viability were determined on a Vi-Cell XR cell viability analyzer (Beckman Coulter, CA, USA) and scRNAseq library preparation was performed using dual single cell mouse kit 5’/TCR according to the manufacturer’s instructions (10x Genomics, CA, USA).

### scRNAseq analysis of STAMP microtumors

Two independent experiments of STAMP microtumors were single cell sequenced using the 10x platform (10x Genomics, CA, USA). The first experiment consisted of four microtumors of each of the following phenotypes: immune infiltrated, immune excluded and immune desert isolated from the same animal to have a shared T cell repertoire across the different tumors. For the second experiment, in addition to repeating the same sampling, we also sequenced tumors derived from a mouse with no adoptively transferred CD3+ T cells and a mouse with only pores (no implantation of tumor cells). For both experiments, paired TCR sequencing was performed. Single-cell RNA-seq data was processed with cellranger count (CellRanger 3.0.2 from 10x Genomics) using a custom reference package based on mouse reference genome GRCm38 and GENCODE^29^ gene models. Data analysis was done in R 3.6.2 and the Seurat package version 3.1.5^33^. Only high quality cells were retained for the posterior analysis, more concretely, we kept the cells with more than 300 hundred genes detected, more than 1000 unique molecule identifiers (UMIs) and less than 10% mitochondrial reads. This filtering step removed most of the doublets and empty cells. From our single cell data, a preliminary clustering was performed using the standard Seurat SCT transform workflow to determine the major cell populations in our samples. The tdTomato positive clusters were analyzed using the standard SCT transform workflow from Seurat. Using the talus plot^34^ we determined that 60 principal components were enough to capture the signal in data. With these retained components, we computed a UMAP embedding and the neighbors for the clustering. Several clustering resolutions were calculated and a directed tree was constructed reflecting the hierarchical relationships of the new clusters upon increasing the resolution^35^. A resolution of 1.5 was considered optimal and the main clusters were identified by means of expression markers of known biology. Markers for each cluster were identified using the Seurat::FindAllMarkers() method with default parameters, comparing all cells in a particular cluster to the rest of cells in the dataset and accessing significantly differential gene expression using Wilcoxon’s rank sum test and FDR correction for multiple testing. We used Seurat’s plotting functionlalities for most plots. Maker heatmaps were generated with the ComplexHeatmap R package^32^ using results from the AverageExpression function of Seurat as input (scaled to relative expression per gene: z-scored per row). Clonotype analyses and integration with Seurat were done using the scRepertoire R package^36^. For the fine structure of the clusters, we were unable to identify a parallel with known T cell subtypes and therefore the clusters were named by their top positive markers. We performed differential gene expression using Seurat::FindMarkers() on each T cell cluster comparing the inflamed to the excluded immune phenotypes using batch as a latent variable and the negative binomial test. We reported the significant genes log Fold Changes values as a z-score scaled matrix using ComplexHeatmap. Clonotypes were called according to their TCR amino acid sequence. Gene Set Enrichment Analysis of the top seven dominant clonotypes was performed on the log Fold Change values from Seurat::FindMarkers() comparing the inflamed to the excluded phenotypes using gseGO() function of the ClusterProfiler R package on the CC Ontology collection.

### 3D in vitro tumoroid cultures/co-cultures

KPP murine pancreatic cancer cells expressing human HER2 and cytosolic GCaMP6 were suspended in 3-dimensional collagen matrices as described in Geraldo et al^37^. In short, a solution of rat-tail collagen I (MilliporeSigma, MA, USA) was brought to a neutral pH on ice and mixed with KPP.hHer2.GCaMP6 cells to a final concentration of 2 mg/mL collagen and 1.0×10^4^ cells. 150 µL of this suspension were added to individual wells of a 8 chambered cover glass (Cellvis, CA, USA). The chambers were incubated at 20-25°C for 15 min then incubated at 37°C with 5% CO_2_ for an additional 15 min. After incubation, 500 µL of complete media containing Gibco RPMI 1640 (Thermo Fisher Scientific, MA, USA) with 10% Gibco fetal calf serum (Thermo Fisher Scientific, MA, USA), 2 mM L-glutamine, and 50 U/mL Gibco penicillin/streptomycin (Thermo Fisher Scientific, MA, USA) was carefully added to each well. Cells were allowed to grow in collagen matrices for 5 days prior to imaging.

3D imaging of OT-I T cells interacting with KPP.hHer2.GCaMP6 tumoroids in collagen matrices was performed on a TiE microscope (Nikon, Japan) with CSU-X1 Spinning Disk (Yokogawa Electric, Japan) and Prime sCMOS camera (Photometrics, AZ, USA). The media in 8 chamber imaging slides containing tumor cell-collagen matrices was replaced with Gibco RPMI 1640 with no phenol red (Thermo Fisher Scientific, MA, USA) with 10% Gibco fetal calf serum (Thermo Fisher Scientific, MA, USA), 2 mM L-glutamine, and 50 U/mL Gibco penicillin/streptomycin (Thermo Fisher Scientific, MA, USA) with 3μM of propidium iodide (Thermo Fisher Scientific, MA, USA). OT-I cells were pre-labeled with Celltrace FarRed (Thermo Fisher Scientific, MA, USA) according to manufacturer’s protocol. FarRed-labeled OT-I cells were added to each chamber containing collagen-suspended KPP.hHer2.GCaMP6 tumoroids and allowed to infiltrate for 2 hours prior to imaging. At the time of imaging, T-cell-dependent bispecific antibody (anti-hHer2::anti-CD3e) was added to the chambers at 500nM final concentration to induce CTL recognition of hHer2-expressing cancer cells. In control conditions, no antibody was added.

### STAMP microtumor correlative imaging of Ca^2+^ influx and T cell infiltration

RAG-2-deficient mice with adoptively transferred tdTomato+ CD3+ T cells bearing mTagBFP2 and GCaMP6-expressing KPP STAMP microtumors were anaesthetized by isoflurane inhalation to effect and imaged daily from 4 days post tumor implantation to 15 days post tumor implantation at the same ear regions. Epifluorescence time-lapse microscopy image series are acquired daily at the same ear regions with a 1.0x Leica PlanApo objective (Leica 10450028) on a Leica M205 FA epifluorescence stereomicroscope every minute for 60-70 min. Image analyses was performed using Imaris software (Bitplane, United Kingdom). Time-lapse image series of individual tumors at 8 days post tumor implantation are semi-automatically segmented and analyzed for Ca^2+^ influx between time points. In addition, tumor sizes, T cell abundances and T cell infiltrations are analyzed. Time-lapse image sequences of individual tumors at 13 days post tumor implantation are semi-automatically segmented and are analyzed for tumor size to determine Day8-to-Day13 tumor growth.

### STAMP microtumor correlative imaging of Ca^2+^ influx and T cell infiltration upon TDB administration

FOXN1-deficient nude mice with adoptively transferred tdTomato+ CD3+ T cells bearing mTagBFP2 and GCaMP6-expressing KPP STAMP microtumors were anaesthetized by isoflurane inhalation to effect and imaged 12 days post tumor implantation. Image series are acquired every 90 seconds for 45 min with a two-photon laser-scanning microscope (Ultima In Vivo Multiphoton Microscopy System, Bruker Technologies, MA, USA) with alternating excitation from dual Ti:sapphire lasers (MaiTai DeepSee, Spectra Physics, MA, USA) tuned to 830 nm and 980 nm, and a 16x numerical aperture 0.8 immersion objective lens (Nikon, Japan). Thereafter, T cell-dependent bispecific antibodies (anti-hHer2::anti-CD3e) are administered intravenously and multiphoton time-lapse microscopy image acquisition is continued at the same region. Time-lapse image series of individual tumors are semi-automatically segmented with Imaris software (Bitplane, United Kingdom) and analyzed for Ca^2+^ influx between time points.

### Image analysis

#### Ca^2+^ influx index in vivo epifluorescence microscopy

An isosurface is created that matches the tumor-associated mtagBFP-2 fluorescence of individual STAMP microtumors. The sum of mtagBFP2 and GCaMP6 fluorescence pixel intensities are calculated for each channel for the tumor isosurface for each time point. The absolute delta of the sum fluorescence intensities between consecutive time points are calculated, averaged for each fluorophore and normalized by the respective mean MFI. The Ca^2+^ influx index is the result of dividing the normalized average delta sum of GCaMP6 intensities by the normalized average delta sum of mtagBFP2 intensities.

#### T cell abundance

An isosurface is created that matches the tumor-associated mTagBFP2 fluorescence of individual STAMP microtumors. The mean fluorescence intensities (MFIs) for the T cell-associated tdTomato fluorescence are determined for the tumor isosurface for each time point. The T cell abundance index is the result of calculating the median of the tdTomato MFIs across all time points.

#### T cell infiltration index

An isosurface is created that matches the tumor-associated mTagBFP2 fluorescence of individual STAMP microtumors. Using the tumor isosurface, two new regions are defined: the tumor center (central 50% of tumor isosurface) and the tumor periphery (area surrounding the tumor that is up to 50µm distance from tumor border). The MFIs for the T cell-associated tdTomato fluorescence are determined for the tumor center and the tumor periphery for each time point and the median of MFIs across all time points is calculated. The T cell infiltration index is the result of the ratio of those medians (center/periphery).

#### Ca^2+^ influx index in vivo 2-photon microscopy

An isosurface is created that matches the tumor-associated mtagBFP-2 fluorescence of individual STAMP microtumors. The sum of mTagBFP2 and GCaMP6 fluorescence pixel intensities are calculated for the tumor isosurface for each time point. The absolute delta of the sum fluorescence intensities between consecutive time points are calculated, averaged for each fluorophore and normalized by the respective mean MFI across all time points. The normalized average delta sum of GCaMp6 intensities is divided by the normalized average delta sum of mTagBFP2 intensities (value1). Also, the average standard deviation of mTagBFP2 and GCaMP6 fluorescence of every pixel of the tumor isosurface is calculated across the time series. The average standard deviation of GCaMP6 is divided by the average standard deviation of mTagBFP2 (value2). The Ca^2+^ influx index (2-photon) is the result of multiplying value1 and value2.

#### Ca^2+^ influx index in vitro spinning disk confocal microscopy

An isosurface is created that matches the tumor-associated GCaMP6 background fluorescence of tumor cells. The median of GCaMP6 MFIs are calculated for the tumor cell isosurfaces for each time point. The absolute delta of the median MFIs between consecutive time points are calculated and averaged. The Ca^2+^ influx index (spinning disk confocal) is the result of dividing the average delta median GCaMP6 MFIs by the mean GCaMP6 MFI across all time points.

#### PI influx index in vitro spinning disk confocal microscopy

An isosurface is created that matches the tumor-associated GCaMP6 background fluorescence of tumor cells. The median of propidium iodide MFIs are calculated for the tumor cell isosurfaces for each time point. The absolute delta of the median MFIs between consecutive time points are calculated and averaged. The propidium iodide influx index is the result of dividing the average delta median propidium iodide MFIs by the mean propidium iodide MFI across all time points.

#### U-net model training

Images of tumor fluorescence are binned and resized to 512×512 using custom FIJI scripts. Binary (2-class) masks are manually generated with 1=Tumor, 0=Background. A TensorFlow U-net model adapted from https://github.com/zhixuhao/unet was trained on a dataset of 595 paired images with masks (70% training and 30% validation) for 7 epochs until the model began to overfit as indicated by the training accuracy exceeding the validation accuracy without improving loss.

#### Image segmentation, tracking

Images of tumor fluorescence are binned and resized to 512×512 using custom FIJI scripts. Initial segmentation guesses are generated by applying the trained U-net using TensorFlow. Custom FIJI scripts are used to un-bin tumor segmentation to restore original size and resolution, enable manual review and editing of all tumor segmentation masks, and manually track tumors through multiple time points. If a tumor is no longer detectable during the course of an experiment, it is designated a complete responder (C.R.). If a tumor decreases from its maximum size by 20% or more, it is designated a partial responder (P.R.). Remaining tumors are designated as stable disease (S.D.) and progressing disease (P.D.).

#### T cell quantification

Custom FIJI scripts are used to identify the centroid and Feret diameter of each tumor ROI and determine the median radial fluorescence profile of the T cell fluorescence channel. Custom Python scripts are used to determine the overall median T cell fluorescence intensity and categorize radial fluorescence profiles as “desert”, “excluded” or “inflamed”. Tumors are classified as “excluded” or “inflamed” using a ratiometric cutoff. If the radial profile within the inner 25% of the tumor is consistently greater than 60% of the maximum fluorescence for that tumor it is designated as inflamed. If the radial profile within the inner 25% of the tumor is consistently less than 40% of the maximum fluorescence for that tumor it is designated as excluded. A tumor is designated a “desert” if the individual tumor’s median T cell fluorescence intensity is less than the 25 percentile of median T cell fluorescence for all tumors measured on the first imaging day and the radial profile does not indicate an excluded pattern as described above. If a tumor shows an excluded profile based on the ratiometric criteria, but the T cell intensity at the core of the tumor (inner 25%) is greater than the median T cell intensity for all inflamed tumors, it is re-classified as inflamed. If a tumor fails to meet the above ratiometric cutoffs, the phenotype determined the previous day is propagated forward until the next definitive classification.

#### Clustering and Markov analysis

Custom python scripts are used to assign an integer value to the T cell phenotype classification with 1=desert, 2=excluded, 3=inflamed. If the mouse is euthanized, the remaining time points are assigned value=0. If the tumor resolves, the remaining time points are assigned value=5. Tumor trajectories are ordered by hierarchical clustering of tumor phenotype lists. Subsequent heatmaps for tumor area, median T cell infiltration and tumor growth are ordered according to this phenotype clustering. Transition state matrices for Markov analysis are generated using custom python scripts. To assess significance, phenotype states are randomized 10 times using Python random.shuffle() and new transition state matrices are calculated.

#### T cell trajectory analysis

Custom python scripts are used to calculate an exclusion ratio where the numerator represents the maximum value of the median radial profile, and the denominator is the median intensity at the core of the tumor (inner 25%). These values are compared between tumors relative to the time-normalized trajectory of each tumor by fitting with a spline (using the SciPy Interpolation sub-package), and then calculating a median value at each time point across all tumors.

#### Multidimension analysis

Data were normalized and standardized (Z-score) before the SPADE tree was built and the EMD scores were calculated.

##### -SPADE trees (Spanning-tree Progression Analysis of Density-normalized Events)

We used the set of four parameters (T cell abundance in the tumor core and tumor periphery, exclusion ratio, and tumor size) to build SPADE trees (SPADE V3.0, http://pengqiu.gatech.edu/software/SPADE/) for n=300. Where n is the number of desired clusters. The total number of tumors (∼11k) were distributed between 300 SPADE tree nodes (details on per node tumor distribution can be found on Extended-Data Fig4.y, bottom, right panel). SPADE views single-object (e.g., cell or tumor) data as a multi-dimensional point cloud and uses topological methods to reveal the geometry of the cloud in 2-dimensional space. Single objects are organized in the 2-dimensional tree structure according to their similarity to each other in the multi-dimensional space. Categorical variables such, e.g. phenotype or response, were converted to the numerical variables that were further used to color the SPADE tree.

##### -EMD scores (Earth Mover’s Distance)

We used the FlowJo V10 (FlowJo, LLC) to run the UMAP analysis (with the default input parameters) on all of the day 6-17 data points simultaneously. The UMAPs were built using a set of four parameters (T cell abundance in the tumor core and tumor periphery, exclusion ratio, and tumor size). We further converted UMAP plots into the density plots for each treatment group (control, anti–TGF-β, anti–PD-L1, and combination of anti–PD-L1 with anti–TGF-β) and calculated the EMD scores^38^ between each treatment group and the group of responders (C.R and P.R). EMD measures the distance between two probability distributions, e.g., the control treatment group and responders’ group.

### Immunophenotypes in imCORE Paired Biopsy trial

Tumor biopsies were obtained from patients enrolled in the imCORE^39^ Paired Biopsy trial (NCT03333655) between January 2018 and March 2020. This study is an ongoing, open-label, multicenter trial initiated in February 2018 and conducted globally including study centers in the United States, France, and Spain. Adult patients with metastatic cancer or hematological malignancies who demonstrated clinical benefit on CIT and had a tumor biopsy both at baseline (pre-treatment/archival) and at progression were eligible for inclusion. CIT included marketed agents (including those targeting CTLA-4, programmed death-ligand 1 [PD-L1] or programmed death-1 [PD-1]) or those administered through participation in a Roche/Genentech CPI clinical trial. Patients with best overall response (per Response Evaluation Criteria in Solid Tumors version 1.1) of complete response (CR), partial response (PR) or stable disease (SD) > 6 months (or > 3 months if enrolled under an earlier protocol version) were eligible. PanCK/CD8 dual staining was performed on histologic sections from baseline and progression formalin-fixed paraffin-embedded tumor samples. Immune phenotypes were determined by a pathologist (Histogenex) using defined criteria^40^.

### Statistics and figure preparation

Statistical analysis was performed by t-test, Mann–Whitney test, one-way ANOVA with Bonferroni’s multiple comparison’s test or log-rank test. P < 0.05 was considered significant. Models in Figs. 1a, 2c, 3g were created using BioRender (https://biorender.com/).

## Supplementary Information

### Supplementary Videos

1) Example of live *in vivo* 2photons imaging of an individual KPP BFP+ STAMP tumor showing the migration of TdTomato+ OTI T cells and WT GFP+ T cell.

2) Epifluorescence imaging of 3 adjacent KPP tumors (magenta) with excluded, inflamed and desert Immune phenotypes, showing T cell migration (cyan) in adjacent tumors (related to Extended Fig.2a)

3) Confocal imaging followed by 3D reconstruction of a whole cleared lung showing KPP-GFP+ tumor foci and TdTomato+ T cells 8 days after i.v transfer. (related to Fig.2b)

4) Confocal live imaging showing low Ca2+ flashes (green) in GCaMP6+ Her2+ KPP organoid, cocultured with T cells (magenta) and propidium Iodide (Red) (related to Fig.3h and Extended Fig.3p)

5) Confocal live imaging showing high Ca2+ flashes (green) in GCaMP6+ Her2+ KPP organoid, cocultured with T cells (magenta)and TDB (aHer2/aCD3) to force tumor cell killing by T cells (positive control). Propidium Iodide entry (Red) reveals tumor cell death. (related to Fig.3h and Extended Fig.3p)

6) *In vivo* 2photons live imaging showing absence of Ca2+ flashes in GCaMP6+ BFP+ Her2+ KPP tumors implanted in Nude mouse (negative control) (Related to Extended Fig.3t)

7) *In vivo* 2photons live imaging showing some Ca2+ flashes in GCaMP6+ BFP+ Her2+ KPP tumors implanted in Nude mouse reconstituted with TdTomato+ T cells (related to Extended Fig.3t)

8) *In vivo* 2photons live imaging showing high level of Ca2+ flashes in GCaMP6+ BFP+ Her2+ KPP tumors implanted in Nude mouse reconstituted with TdTomato+ T cells and treated with TDB (aHer2/aCD3) to force tumor cell killing by T cells (positive control) (related to Extended Fig.3t)

9) Epifluorescence live imaging of an Immune Inflamed tumor showing high Ca2+ flashes in GCaMP6+ KPP tumor cells implanted in Nude mouse reconstituted with TdTomato+ T cells (related to Fig3j)

10) Epifluorescence live imaging of an Immune excluded tumor showing low Ca2+ flashes (green) in GCaMP6+ KPP tumor cells implanted in Nude mouse reconstituted with TdTomato+ T cells (related to Fig.3j)

### Supplementary Tables

1) Markov Probabilities of Transition between Immune Phenotypes (related to Fig.4h and Extended Fig.4i-j)

2) Raw measurements of 14300 KPP-GFP STAMP tumors implanted into RAG2-/-mice reconstituted with TdTomato+ T cells and treated with CIT one Day after tumor implantation (related to Fig.4)

3) Clinical summary (related to Fig.4q)

## Acknowledgements

We thank members of C. Moussion, S.J. Turley, and I. Mellman laboratories for advice, discussions and reagents; B. Hough & R. Asuncion, for animal husbandry; R. Garcia-Gonzalez, J. Yamada & E. Chua for veterinary care; the Genentech FACS group for technical assistance; HyperVoxel, Johannes Schoeneberg and Gautham Raghupathi for the support with Image processing and machine learning and the Genentech postdoctoral program for support. This study was funded by Genentech/Roche.

## Author contributions

G.O.M and M.B. designed and performed the experiments, analyzed and interpreted the data, C.B.C developed the high content image analysis pipeline to analyze tumors features over-time, analyzed and interpreted the STAMP CIT timeseries experiments, J.P.J analyzed and interpreted NGS data, D.O analyzed and interpreted the STAMP CIT timeseries experiments, A.T.R., J.D, S.M, A.Y performed experiments; S.T edited the manuscript, Z.M supervised sequencing, T.W participated in single cell analysis, R.P. supervised the collection of clinical samples; I.M discussed the data and edited the manuscript; Y.W and S.M shared clinical trial data, C.G-R, C.E.A, D.S collected patient samples for imCORE studies, C.M. conceived and supervised the study, interpreted the data and wrote the manuscript; and all authors read and approved the final article.

