## Supplementary Figures for "Surveillance of *in situ* tumor arrays reveals early environmental control of cancer immunity"

### Extended-Data Fig. 1 | Implantation and growth analysis of clonally-derived tumors

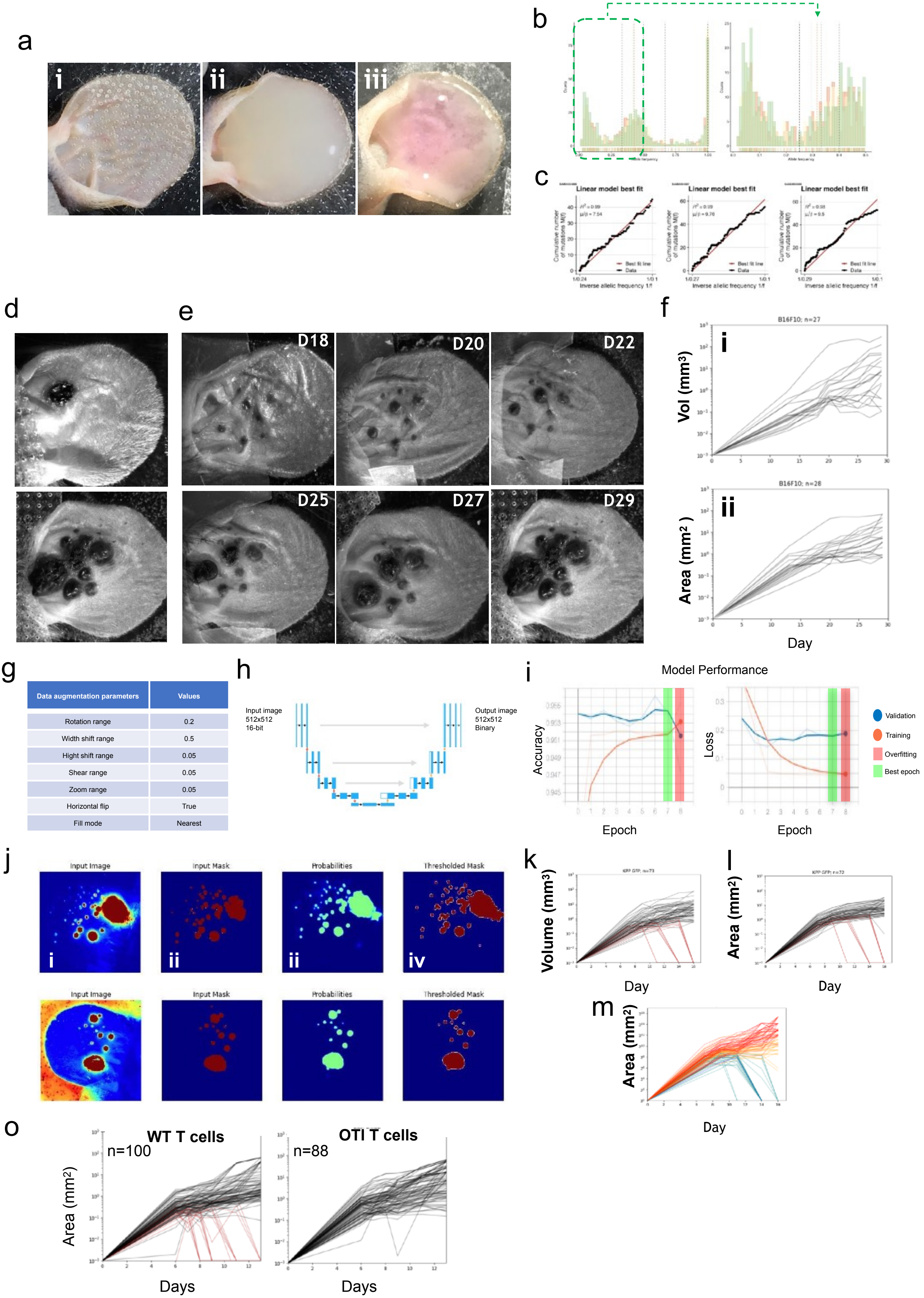


| Data augmentation parameters | Values |
|---|---|
| Rotation range | 0.2 |
| Width shift range | 0.5 |
| Hight shift range | 0.05 |
| Shear range | 0.05 |
| Zoom range | 0.05 |
| Horizontal flip | True |
| Fill mode | Nearest |

**Extended Data Fig. 1 | Implantation and growth analysis of clonally-derived tumors**
**a,** STAMP (<u>s</u>kin <u>t</u>umor <u>a</u>rray via <u>m</u>icroporation) tumor implantation. (i) Representative image of ears after P.L.E.A.S.E.® Laser microporation, (ii) microporated ears covered with tumor cell suspension, (iii) microporated ear seeded with tumor cells and covered with Matrigel. **b,** Variant allele frequency (VAF) histogram of the KPP-EGFP cell line highlighting it's clonality. The dashed colored lines indicate the median VAF for each distribution. Black dashed lines stand for homozygous and the heterozygous peaks as well as 1/3 and 2/3 copy number peaks. **c,** Neutrality test for each VAF distribution. **d,** Representative B16F10 tumor arrays 29 days post-implantation of primary/native tumor cell line (upper panel) and in-vivo passaged secondary cell line (lower panel) to show improvement of engraftment **e,** Representative time course image series of secondary B16F10 microtumors as described in (d). **f,** Manual analyses of growth kinetics of individual B16F10 STAMP tumor using ellipsoid formula to calculate volumes over time (i) and using tumor perimeter to calculate area over time (ii). n=27 tumors, 2 animals pooled. **g-j**, Development of the high content image analysis pipeline: **g.** Data augmentation parameters for training data (416 frames) and validation data (119 frames). **h**, Single class segmentation process from input epifluorescence image to output tumor mask and overview of the U-net architecture. **i**, Performance of models during training for tumor segmentation. **j,** Two representative tumor segmentations (upper and lower panel) performed on validation images. Input images (i), manually generated classification mask (ii), features extracted after the penultimate upsampling step (iii), output segmentation mask (iv). **k-m,** Validation of the high content image analysis pipeline. **k,** Manual analyses of growth kinetics of individual KPP-EGFP tumor volumes ($\mu m^3$) using the ellipsoid formula. n=72 tumors, 3 animals pooled. **l,** Manual analyses of growth kinetics of individual tumor areas ($\mu m^2$) using tumor perimeter. n=72 tumors, 3 animals pooled. **m,** Automated analyses of growth kinetics of individual tumor areas ($\mu m^2$) n=72 tumors, 3 animals pooled **o,** Automated analysis of growth kinetics of individual tumor area (mm^2) for experiment described in Fig 1.i. Mice reconstituted by adoptive transfer of CD3+ T cells from either C57BL/6J WT (left panel) or OT-1+ CD4-cre tdTomato+ (right panel) n≧88 tumors. Red lines indicate tumors that are rejected, gray lines indicate tumors that persist.

### Extended-Data Fig. 2 |
### Heterogeneity and clinical relevance of mouse STAMP tumor-Immune phenotypes

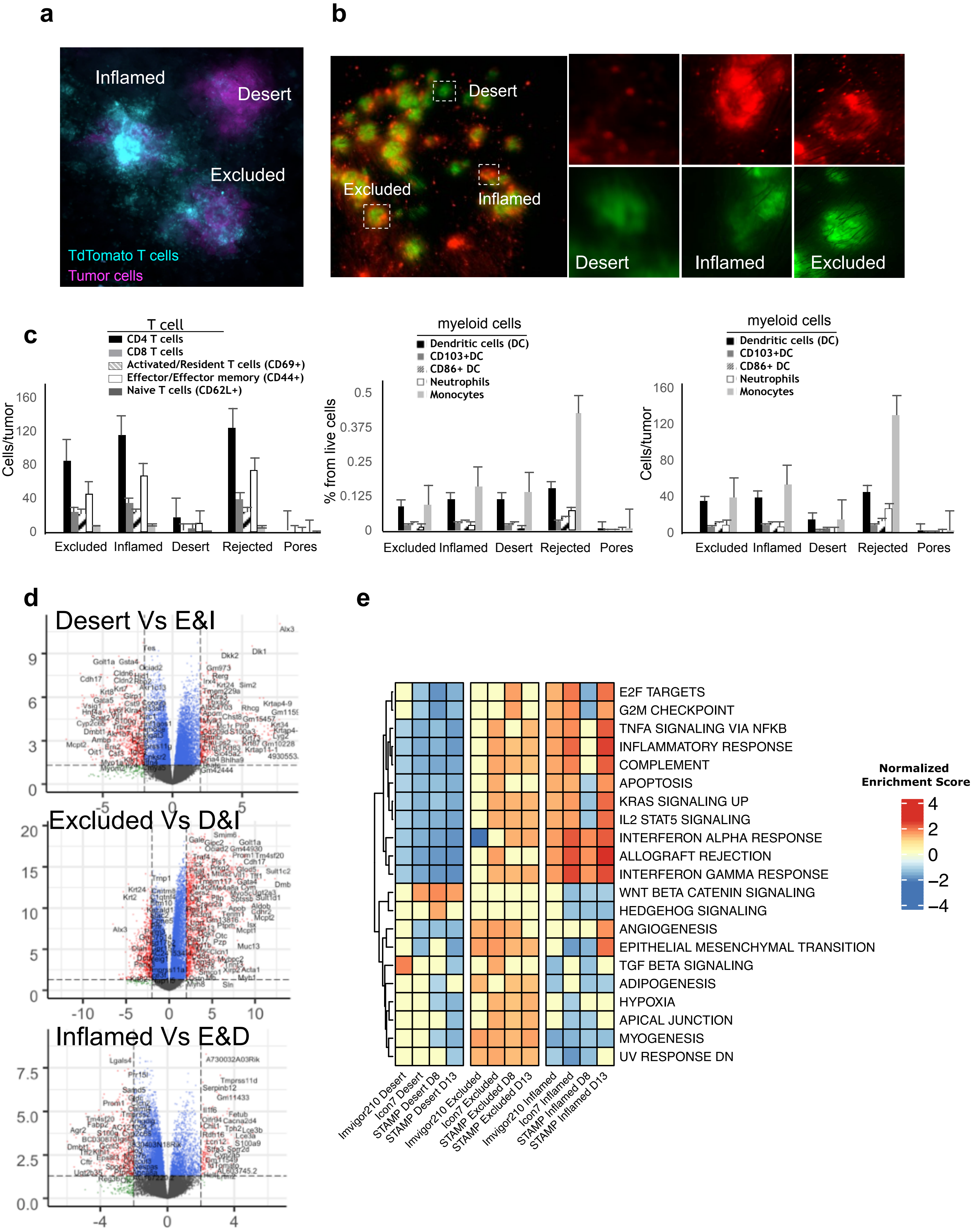

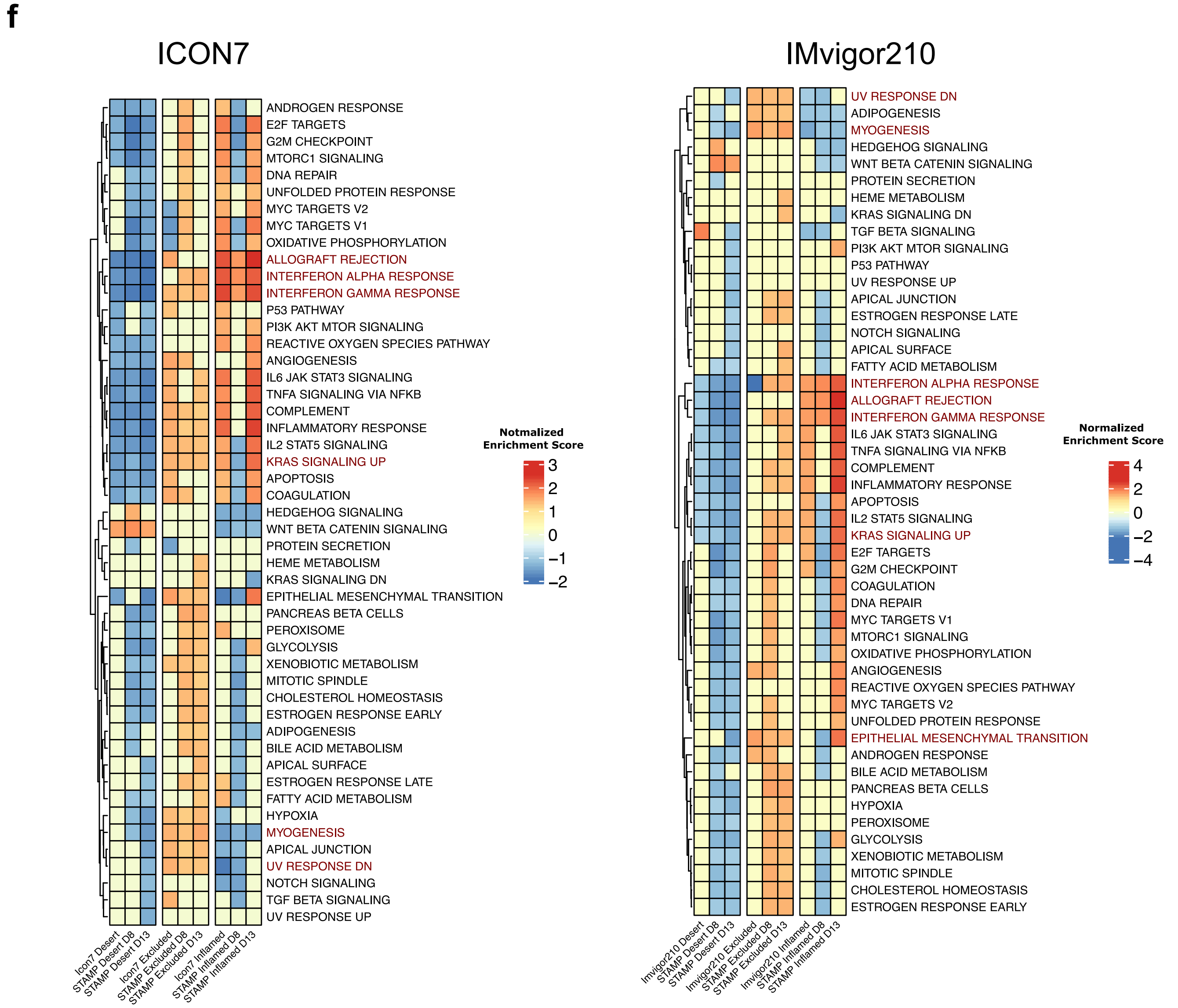


**Extended Data Fig. 2 | Heterogeneity and clinical relevance of mouse STAMP tumor-Immune phenotypes**
**a,** Representative STAMP images (movie in supplement) of KPP-EGFP on ears of C57BL/6J Foxn1-Nude mice at 8 days post tumor implantation, reconstituted by adoptive transfer of CD3+ T cells from C57BL/6J WT CD4-cre tdTomato+ mice (T cells – cyan, KPP-EGFP – magenta). Individual tumors with differential immune phenotypes are in close proximity to each other. n=5 animals. **b,** Representative STAMP of KPP-EGFP on shaved abdominal skin of C57BL/6J at 8 days post tumor implantation, RAG-2-deficient mice reconstituted by adoptive transfer of CD3+ T cells from C57BL/6J WT CD4-cre tdTomato+ mice (T cells – red, KPP-EGFP – green). Left panel represents an overview of the entire abdominal tumor array, right panels are enlarged images of individual tumors with diverse immune phenotypes: inflamed, excluded, desert, resolved. n=3 animals. **c,** Flow cytometry-based immune cell profiling of individual tumor biopsies pooled by immune phenotype at 10 days post tumor implantation. n=6 animals, ≧ 3 pools of tumors **d,** Volcano plots of differentially expressed genes (q-value< 0.05) between individual tumors of three different immune phenotypes (inflamed, desert, excluded) with an absolute logFC greater than 2. **e,** Heat map comparing the normalized enrichment scores for most conserved pathways that are significantly enriched between different immune phenotypes from ICON7 (ovarian), IMvigor210 (bladder), IMvigor211 (bladder) clinical trials tumor samples, and STAMP individual tumors at day8 and at day13. **f,** Heat map comparing the normalized enrichment scores for all pathways that show significance in at least one comparison of the different immune phenotypes of each individual clinical trial and STAMP tumors.

**Extended-Data Fig. 3 |**

**Immune phenotype determines T cell function regardless of T cell clonotype**

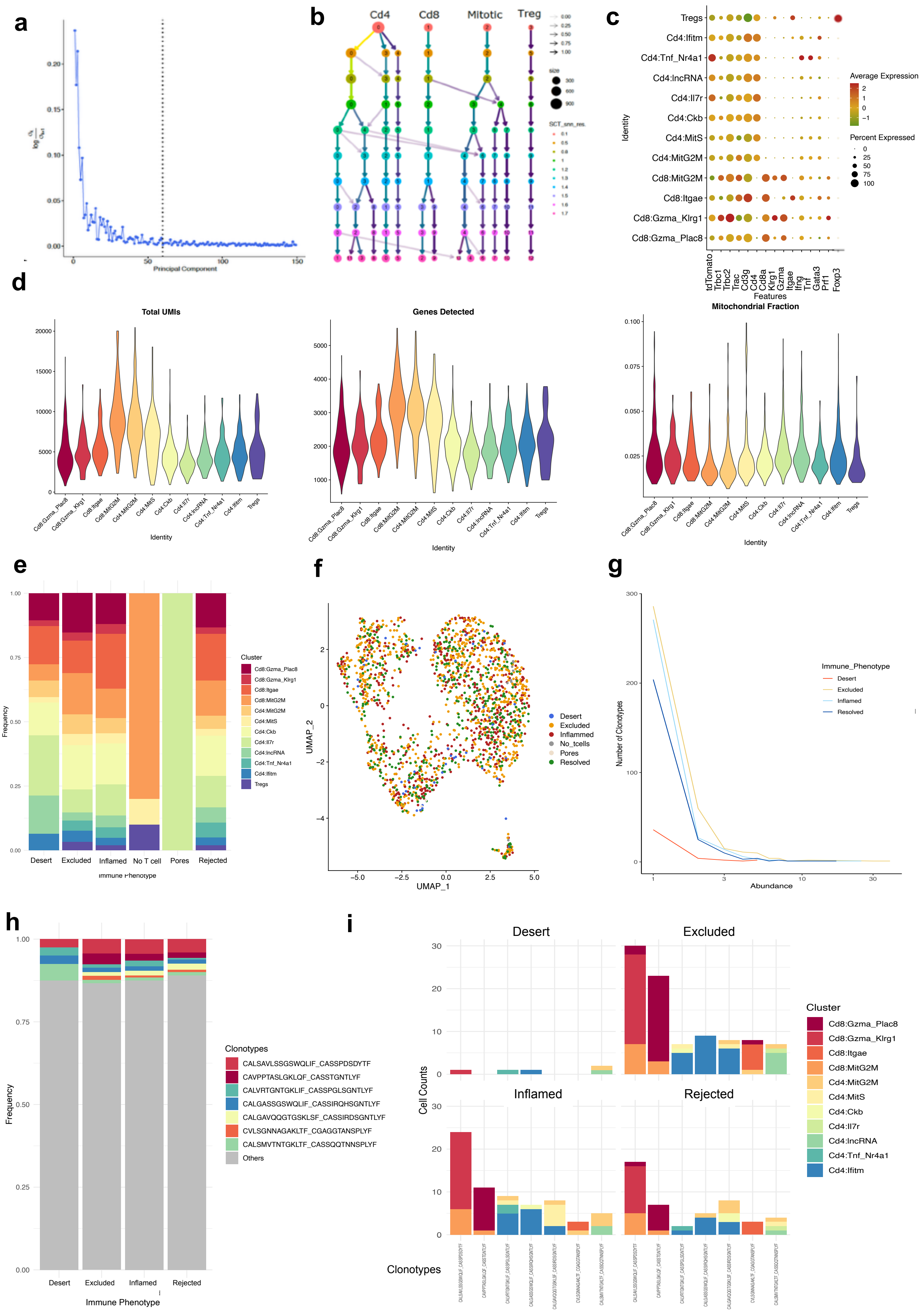

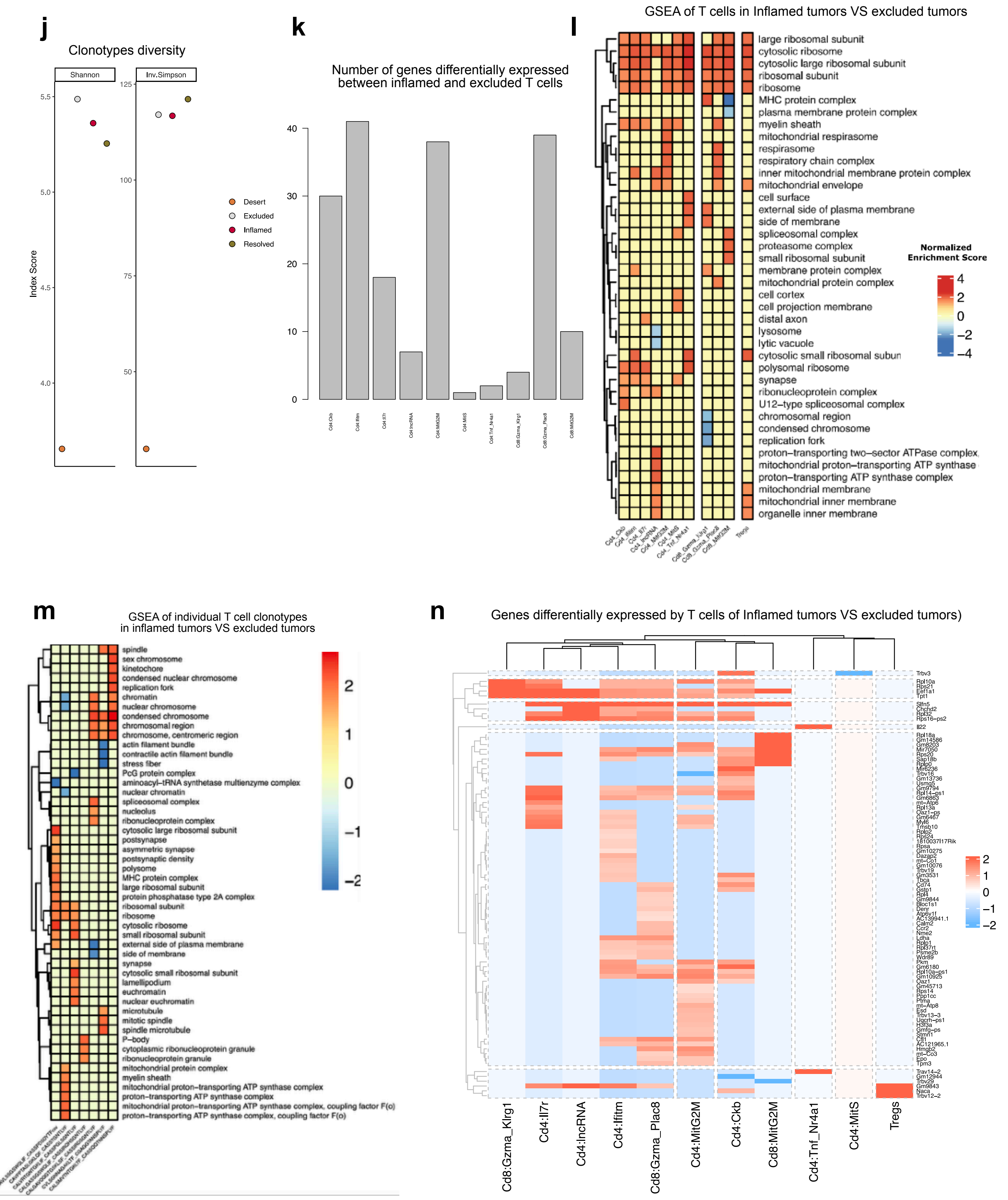
j
Clonotypes diversity
Shannon
Inv.Simpson
Index Score
Desert
Excluded
Inflamed
Resolved
k
Number of genes differentially expressed
between inflamed and excluded T cells
l
GSEA of T cells in Inflamed tumors VS excluded tumors
large ribosomal subunit
cytosolic ribosome
cytosolic large ribosomal subunit
ribosomal subunit
ribosome
MHC protein complex
plasma membrane protein complex
myelin sheath
mitochondrial respirasome
respirasome
respiratory chain complex
inner mitochondrial membrane protein complex
mitochondrial envelope
cell surface
external side of plasma membrane
side of membrane
spliceosomal complex
proteasome complex
small ribosomal subunit
membrane protein complex
mitochondrial protein complex
cell cortex
cell projection membrane
distal axon
lysosome
lytic vacuole
cytosolic small ribosomal subun
polysomal ribosome
synapse
ribonucleoprotein complex
U12−type spliceosomal complex
chromosomal region
condensed chromosome
replication fork
proton−transporting two−sector ATPase complex
mitochondrial proton−transporting ATP synthase
proton−transporting ATP synthase complex
mitochondrial membrane
mitochondrial inner membrane
organelle inner membrane
Normalized
Enrichment Score
Tregs
m
GSEA of individual T cell clonotypes
in inflamed tumors VS excluded tumors
spindle
sex chromosome
kinetochore
condensed nuclear chromosome
replication fork
chromatin
nuclear chromosome
condensed chromosome
chromosomal region
chromosome, centromeric region
actin filament bundle
contractile actin filament bundle
stress fiber
PcG protein complex
aminoacyl−tRNA synthetase multienzyme complex
nuclear chromatin
spliceosomal complex
nucleolus
ribonucleoprotein complex
cytosolic large ribosomal subunit
postsynapse
asymmetric synapse
postsynaptic density
polysome
MHC protein complex
large ribosomal subunit
protein phosphatase type 2A complex
ribosomal subunit
ribosome
cytosolic ribosome
small ribosomal subunit
external side of plasma membrane
side of membrane
synapse
cytosolic small ribosomal subunit
lamellipodium
euchromatin
nuclear euchromatin
microtubule
mitotic spindle
spindle microtubule
P−body
cytoplasmic ribonucleoprotein granule
ribonucleoprotein granule
mitochondrial protein complex
myelin sheath
mitochondrial proton−transporting ATP synthase complex
proton−transporting ATP synthase complex
mitochondrial proton−transporting ATP synthase complex, coupling factor F(o)
proton−transporting ATP synthase complex, coupling factor F(o)
n
Genes differentially expressed by T cells of Inflamed tumors VS excluded tumors)
Cd8:Gzma_Klrg1
Cd4:Il7r
Cd4:lncRNA
Cd4:Ifitm
Cd8:Gzma_Plac8
Cd4:MitG2M
Cd4:Ckb
Cd8:MitG2M
Cd4:Tnf_Nr4a1
Cd4:MitS
Tregs

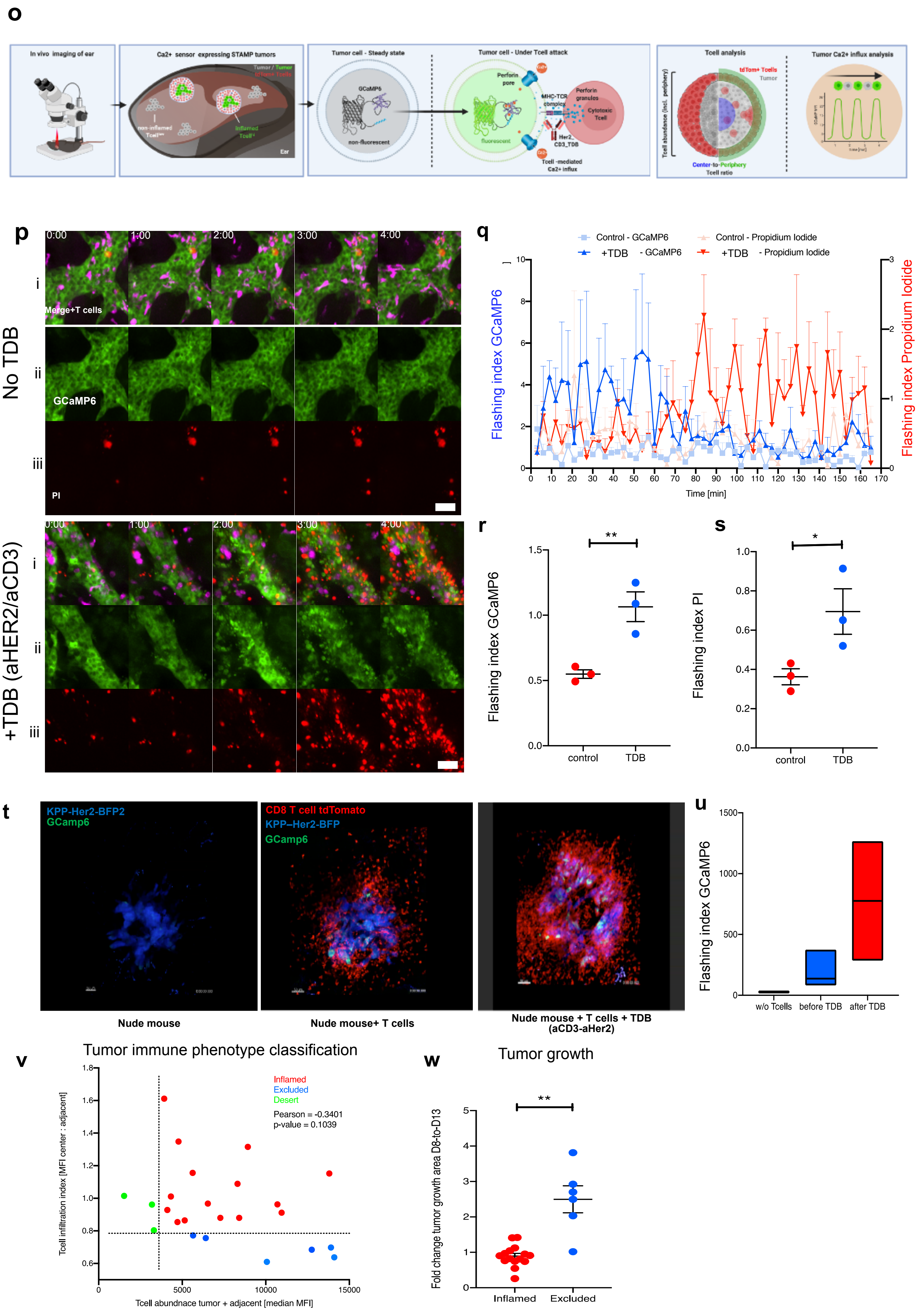
o
In vivo imaging of ear
Ca2+ sensor expressing STAMP tumors
Tumor / Tumor
tdTom+ Tcells
non-inflamed Tcell
inflamed Tcell
Ear
Tumor cell - Steady state
GCaMP6
non-fluorescent
Tumor cell - Under Tcell attack
Perforin pore
fluorescent
MHC-TCR complex
Perforin granules
Cytotoxic Tcell
Her2_CD3_TDB
Tcell -mediated Ca2+ influx
Tcell analysis
tdTom+ Tcells
Tumor
Tcell abundance (incl. periphery)
Center-to-Periphery Tcell ratio
Tumor Ca2+ influx analysis
p
No TDB
0:00
1:00
2:00
3:00
4:00
i
Merge+T cells
ii
GCaMP6
iii
PI
+TDB (aHER2/aCD3)
q
Control - GCaMP6
+TDB - GCaMP6
Control - Propidium Iodide
+TDB - Propidium Iodide
Flashing index GCaMP6
Flashing index Propidium Iodide
Time [min]
r
Flashing index GCaMP6
control
TDB
**
s
Flashing index PI
control
TDB
*
t
KPP-Her2-BFP2
GCamp6
CD8 T cell tdTomato
KPP–Her2-BFP
GCamp6
Nude mouse
Nude mouse+ T cells
Nude mouse + T cells + TDB (aCD3-aHer2)
u
Flashing index GCaMP6
w/o Tcells
before TDB
after TDB
v
Tumor immune phenotype classification
Inflamed
Excluded
Desert
Pearson = -0.3401
p-value = 0.1039
Tcell infiltration index [MFI center : adjacent]
Tcell abundnace tumor + adjacent [median MFI]
w
Tumor growth
Fold change tumor growth area D8-to-D13
Inflamed
Excluded
**

**Extended Data Fig. 3 | Immune phenotype determines T cell function regardless of T cell clonotype**

**a,** The talus plot indicates that 60 principal components are sufficient to capture the signal of the single cells data. **b,** Cluster tree with different clustering resolutions, 1.5 is sufficient to obtain the most robust clustering. **c,** Dot plot for the marker genes of the main T cell clusters. **d,** Quality control measures by cluster. **e,** Relative abundance for each of the subclusters separated by immune phenotypes. **f,** UMAP colored by tumor immune phenotypes **g,** Rank abundance plots of T cell clonotypes. **h,** Relative abundance of T cell clonotypes, **i,** cell number of each T cell clonotype with cluster identification **j,** TCR clonotype diversity indices for each immune phenotype. **k,** Number of genes differentially expressed between Inflamed and Excluded T cell clusters **l,** GSEA for each T cell subcluster of Inflamed VS excluded tumors, top ten pathways ordered by p-value (descending) **m,** GSEA performed on individual T cell clonotypes seating in inflamed versus excluded tumors **n,** Genes differentially expressed between Inflamed and excluded T cells per clusters (red=high in Inflamed tumors, blue=high in excluded tumors). **o-u,** Validation of GCaMP6 expressing tumor cells as a tool to monitor tumor cell attack by T cells in vitro and in vivo. **o,** Schematic workflow of the live *in vivo* imaging and quantification of Tumor cell attacks by T cells in STAMP tumor arrays. Ears of Rag2 KO mice are imaged by epifluorescence microscopy on day 8 after the seeding of GCaMP6 expressing tumor cells and adoptive transfer of tdTomato+ Tcells. Cytotoxic T cells create Ca2+-permeable perforin pores in the tumor cell membrane. Ca2+ influx activates GCaMP6 fluorescence. Schematic of Tcell abundance, Tcell infiltration and Ca2+ flux analysis. **p,** Representative *in vitro* time-lapse image series of GCaMP6-Her2-expressing tumor organoids cultured with TdTomato T cells in the absence (upper panel) or presence (lower panel) of aHer2-aCD3 Tcell-dependent bispecific antibody to force tumor cell killing by T cells (TDB). (i) Composite of GCaMP6 (green), propidium iodide (PI-red) and Tcell fluorescence (magenta), (ii) single channel of GCaMP6 fluorescence, (iii) single channel of PI fluorescence. n=3. **q,** Average Ca2+ influx indices (blue) and PI death indices (red) plotted against the time (min) in the absence (control, light color) or presence (TDB, dark color) of TDB as described in o. n=3. **r,** Ca2+ influx indices in the absence (control) or presence (TDB) of TDB as described in p. n=3. *p-value ≦ 0.01, unpaired one-sided T-test **s,** PI death indices in the absence (control) or presence (TDB) of TDB as described in p. n=3. *p-value ≦ 0.05, unpaired one-sided T-test. **t,** Representative *in vivo* time-lapse images of GCaMP6-Her2 expressing STAMP tumors in Rag2KO mice on Day12 after tumor cell seeding. Animals did not (left panel) or did (middle and right panel) receive an adoptive transfer of tdTomato+ T cells. Mice that received adoptive Tcell transfer were imaged before (middle panel) and after (right panel) I.V. administration of TDB to force Tumor/T cell interaction and tumor killing (positive control). GCaMP6: green, KPP tumor cells: blue, and tdTomato Tcells: red. n≧2. **u.** Ca2+ influx indices in the absence of Tcells (w/o Tcells- negative control), before TDB or after TDB (positive control) as described in I. n≧2. **v,** Systematic tumor immune phenotype classification for inflamed, excluded and desert tumors as described in (Fig3.j-n). **w,** Tumor growth fold change Day8-to-Day13 of inflamed and excluded tumors as described in (Fig3.j-n). n≧6. **** p-value ≦ 0.0001, unpaired T-test with Welch’s correction.

**Extended data Fig. 4 |**
**Early transition to an inflamed phenotype predicts response to Immunotherapy**

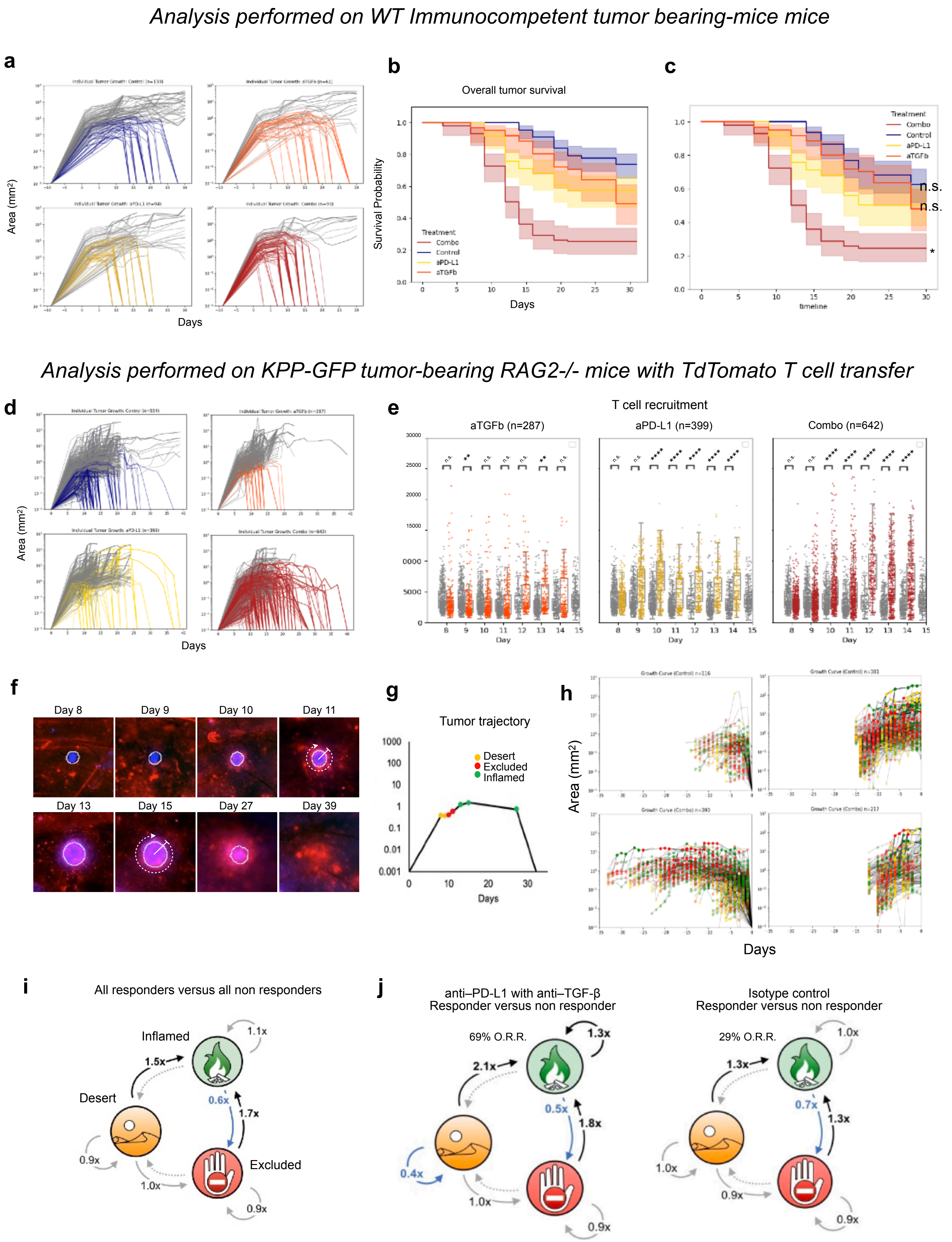

k

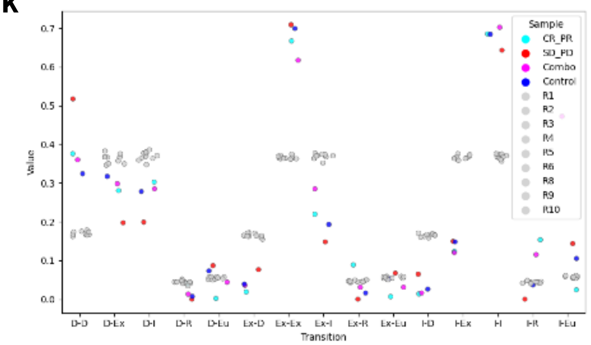


l

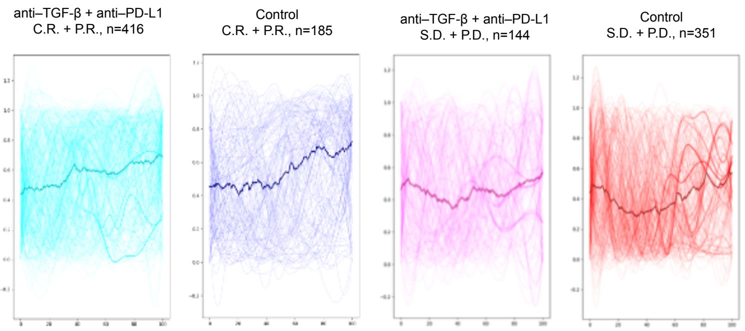


m

*Multidimensional UMAP analysis and Earth Moving Distance calculation*

*KPP-GFP tumor-bearing RAG2-/- mice with TdTomato T cell transfer*

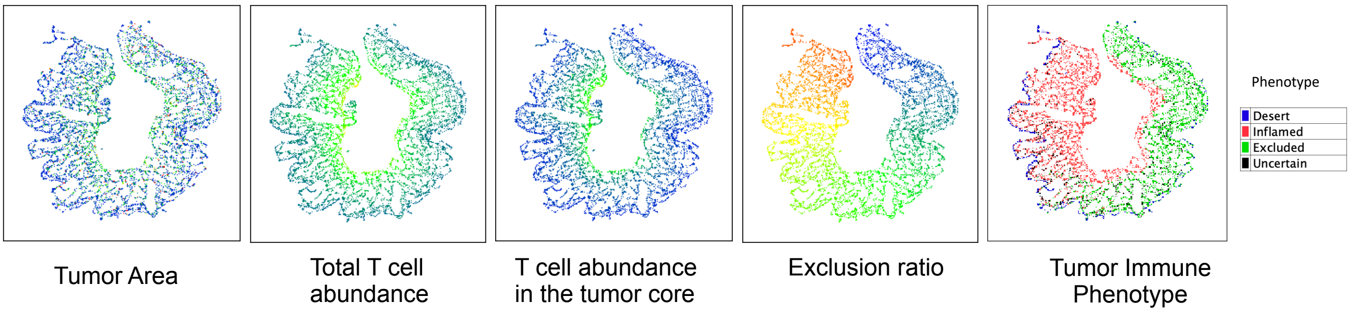


n

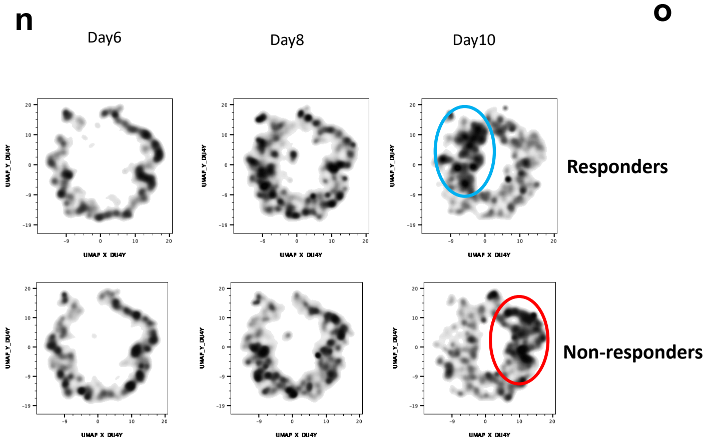


o

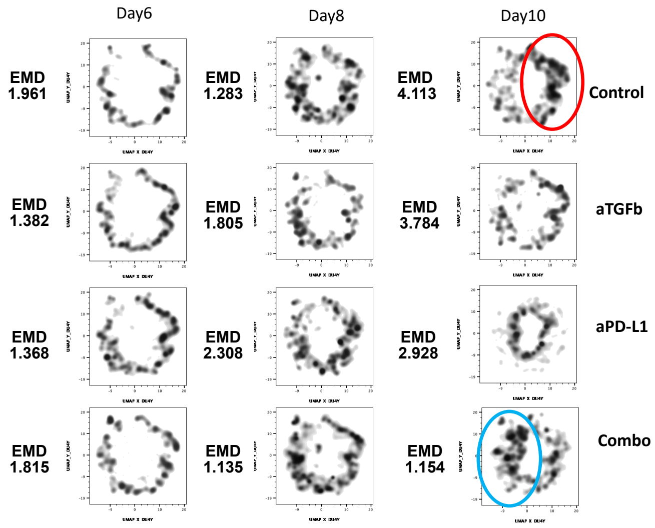


p

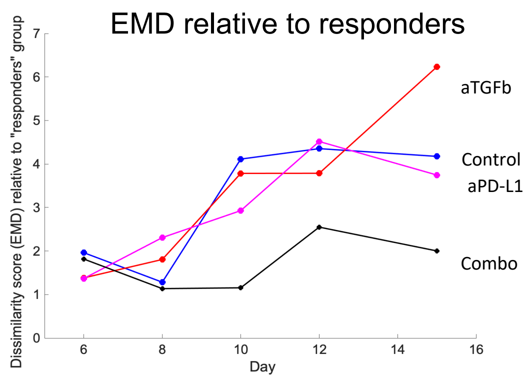


q

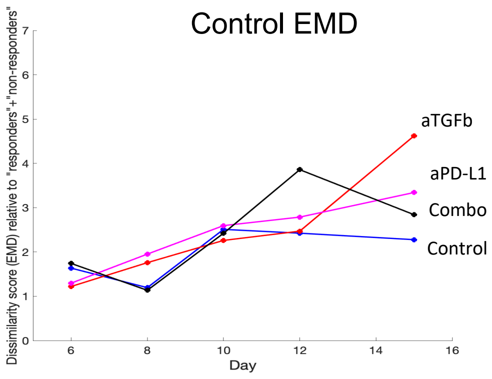

*Multidimensional SPADE tree representation of 11K tumors*
*KPP-GFP tumor-bearing RAG2-/- mice with TdTomato T cell transfer*

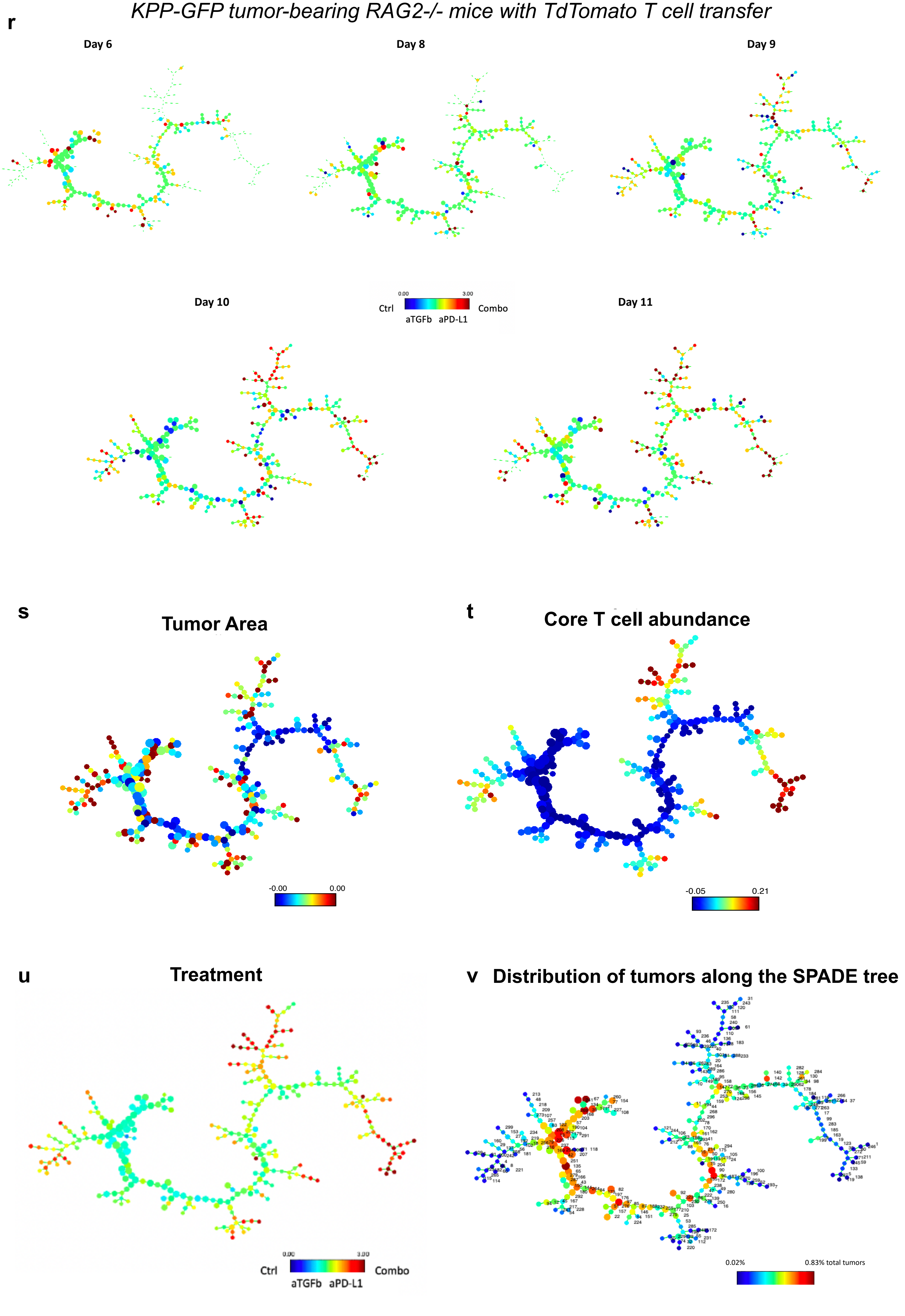

**Fig. 4 | Early transition to an inflamed phenotype predicts response to Immunotherapy**

**a,** Individual tumor growth kinetics (mm^2) of KPP-GFP STAMP implanted in WT animals treated at day 5 post-implantation with isotype control antibodies, anti–PD-L1, anti–TGF-β or a combination of anti–PD-L1 with anti–TGF-β. n=61-130 tumors, 3-4 animals/group. Colored lines indicate tumors that are rejected, gray lines indicate tumors that persist.. **b,** Kaplan Meier survival curves of individual tumors as described in a. Shaded area=95% confidence interval, log-rank test p-value n.s. >0.5, ****<0.0001. **c,** Left censored Kaplan Meier survival curves of individual microtumors as described in a. **d**,Individual tumor growth kinetics (mm^2) of KPP-BFP STAMP implanted in Rag2 KO animals reconstituted with tdTomato T cells and and treated at day 1 post-implantation with isotype control antibodies, anti–TGF-β, anti–PD-L1, or a combination of anti–PD-L1 with anti–TGF-β. n≧287 tumors per group and 10-12 animals per group. **e,** Total T cell abundance (tdTomato MFI per tumor) for individual microtumors as described in d. Isotype control antibodies (gray dots showed as reference), anti–TGF-β (orange dots), anti–PD-L1 (yellow dots), or a combination of anti–PD-L1 with anti–TGF-β (red dots). **f**, Kinetics of T cells (red) and tumor cell (blue) over time for an individual tumor **g,** overlay of the automated classification of immune phenotypes with the individual tumor growth curve for the example in (f). **h,** Tumor growth and immune phenotype trajectory of rejected tumor (left panels) aligned at rejection time and progressing tumors (right panels) aligned at takedown time, (desert-yellow, inflamed-green, excluded-red) as described in d. (n=287-642 microtumors, 10-12 animals/group). **i**, Markov chain showing the difference between transition matrices for all responders (C.R. + P.R.) versus all non responder tumors (S.D. + P.D.) or **j.** broken down by treatment arm. combo treated group (left) or control group (right). **k,** Transition matrix for data represented in Fig. 4h compared to 10 equivalent randomly simulated datasets **l,** Exclusion ratio over normalized tumor trajectory for individual tumors and average as described in Fig. 4k. Combo C.R. + P.R. in cyan, combo S.D. + P.D. in magenta. Control C.R. + P.R. in blue, control S.D. + P.D. in red. **m,** UMAP plots of 11K tumors for day 6-17 are color coded based on Area, Abundance, Core, Ratio and Phenotype parameter values, where blue color represents smaller parameter values and red color represents higher parameters values. FlowJo V10 (FlowJo, LLC) is used to run the UMAP analysis on all of the data points simultaneously. **n,** density distributions in the UMAP coordinates broken down by responders and non-responders groups at day 6, 8 and 10. **o**, For day 6, 8, 10 the the UMAP density plots were broken down by treatment groups and the EMD dissimilarity scores were calculated between each treatment group and the group of responders (C.R and P.R). **p**, Time course of the EMD scores between each treatment group and the group of responders and **q,** the control analysis of EMD scores between each treatment group and all tumors (responders+non-responders).

**r,** Evolution of the SPADE tree over the course of five days. The SPADE tree is color coded according to the treatment group. **s-v**, SPADE tree representing all of the day 6-17 data points simultaneously. SPADE trees were color-coded according to Tumor Area (**s**), T cell abundance in the core of the tumor **(t)**, treatment type (**u)** and distribution of tumors along the 300 nodes of the Spade Tree (**v)**
